# EGFR^+^ lung adenocarcinomas coopt alveolar macrophage metabolism and function to support EGFR signaling and growth

**DOI:** 10.1101/2023.04.15.536974

**Authors:** Alexandra Kuhlmann-Hogan, Thekla Cordes, Ziyan Xu, Kacie A. Traina, Camila Robles-Oteíza, Deborah Ayeni, Elizabeth M. Kwong, Stellar R. Levy, Mathew Nobari, George Z. Cheng, Reuben Shaw, Sandra L. Leibel, Christian M. Metallo, Katerina Politi, Susan M. Kaech

**Author notes:** Corresponding authors: Susan M. Kaech, PhD, NOMIS Center for Immunobiology and Microbial Pathogenesis Salk Institute, Katerina Politi, PhD, Departments of Pathology and Internal Medicine (Section of Medical Oncology) Yale Cancer Center, Yale University School of Medicine.

## Abstract

The limited efficacy of currently approved immunotherapies in EGFR-mutant lung adenocarcinoma (LUAD) underscores the need to better understand mechanisms governing local immunosuppression. Elevated surfactant and GM-CSF secretion from the transformed epithelium induces tumor-associated alveolar macrophages (TA-AM) to proliferate and support tumor growth by rewiring inflammatory functions and lipid metabolism. TA-AM properties are driven by increased GM-CSF—PPARγ signaling and inhibition of airway GM-CSF or PPARγ in TA-AMs suppresses cholesterol efflux to tumor cells, which impairs EGFR phosphorylation and restrains LUAD progression. In the absence of TA-AM metabolic support, LUAD cells compensate by increasing cholesterol synthesis, and blocking PPARγ in TA-AMs simultaneous with statin therapy further suppresses tumor progression and increases T cell effector functions. These results reveal new therapeutic combinations for immunotherapy resistant EGFR-mutant LUADs and demonstrate how such cancer cells can metabolically co-opt TA-AMs through GM-CSF—PPARγ signaling to provide nutrients that promote oncogenic signaling and growth.

## Introduction

Lung cancer remains the leading cause of cancer-related mortality worldwide with a 5-year survival rate of less than 25%. Lung adenocarcinoma (LUAD), the most common histopathological subset of lung cancer, arises following transformation of lung epithelial cells, most commonly type II pneumocytes (AT2), which perform secretory and regenerative functions in the alveolus (1). Molecular studies of LUADs have identified genetic alterations in *KRAS*, the epidermal growth factor receptor (*EGFR*) and anaplastic lymphoma kinase (*ALK*) as major oncogenic drivers of LUAD (2). While LUADs with *EGFR* mutations and *ALK* rearrangements typically respond well to oncogene-targeted therapies, resistance inevitably emerges within 12-18 months (3). Immunotherapies, though more durable, are efficacious in a smaller percentage of LUAD patients but are not generally effective in tumors driven by mutant *EGFR*. In particular, high neoantigen burden, expression of immune checkpoint molecules, and T cell infiltration correlate with immune checkpoint blockade (ICB) response rates, but since EGFR^+^ LUADs lack these characteristics, EGFR^+^ tumors are mostly unresponsive to ICB (4,5). Thus, a need exists for alternate therapeutic approaches that can harness patients’ immune systems to fight non-T cell inflamed tumors, like EGFR^+^ LUADs.

Tumor associated macrophages (TAMs) pervade even the most poorly T cell-infiltrated tumors. Patients with a high density of macrophages (6–8) or elevated levels of the chemokines that recruit them (9,10) have worse outcomes and preclinical studies have shown that inhibiting myeloid cell recruitment can be effective (9,11–13) evidencing a generally tumor supportive role for TAMs. However, TAMs, like all macrophages, are plastic and function across a spectrum of opposing functional axes that can support or hinder tumorigenesis—i.e. inflammatory *vs.* anti-inflammatory, tissue repair *vs.* fibrosis, nutrient competition vs support. TAMs can secrete pro-angiogenic factors which facilitate the mobilization of nutrients and oxygen into tumors (14) as well as growth factors to directly promote oncogenic signaling or immune cell infiltration and activation in tumors (15,16). Inflammatory cytokine secretion by TAMs has the potential to evoke an anti-tumor immune response but can also be maladaptive and drive tumor progression (17) and T cell exhaustion (18). The preclinical success of TAM repolarization by agonistic CD40 underscores the potential of toggling TAM functional states therapeutically for patients whose tumors respond poorly to ICB (19,20). Thus, there is a need to better delineate the tumor-supportive functions of TAMs to uncover novel vulnerabilities, particularly in cold tumors, and develop new therapeutic interventions targeting myeloid populations (21–23).

TAMs are comprised of both recently recruited monocyte-derived macrophages and tissue resident macrophages (TRMs). TRMs evolved to meet the functional demands of the unique microenvironmental needs of their ‘client’ parenchymal cells (24–26) and, on the one hand, are critical to tissue homeostasis (27). Such functions include synaptic pruning from microglia (28); bone resorption from osteoclasts (29), promoting β-cell homeostasis by islet cell macrophages (30,31) and surfactant metabolism by alveolar macrophages (AMs) (32). However, on the other hand, TRMs can also promote tumorigenesis in each of these tissues (33–37), but how their tissue-specialized functions contribute to tumorigenesis remains largely unexplored. For example, in the bone, metastatic cancer cells hijack osteoclasts to induce bone resorption to establish a tumor-supportive niche (35). Metabolic adaptations of TAMs have been well characterized in tumors (38), but there is a need to better understand more specifically how their metabolic status is influenced by their tissue of origin and the metabolic demands of their transformed ‘client’ cells. We hypothesize that tumor cell co-option of homeostatic TRM relationships, including their metabolic relationships, can expose targetable vulnerabilities for cancer treatment.

In the lung, AMs play a pivotal role in maintaining metabolic homeostasis by regulating the surfactant pool. Lung surfactant is a complex mixture of lipids and proteins secreted by AT2 cells to facilitate gas exchange. AMs and AT2 cells cooperate to tightly regulate surfactant composition to maintain its biophysical properties, while heightened lipid metabolism in AMs also feeds into their critical role in maintaining airway tolerance. In this study we identify a metabolic and immunologic feedback loop between tumor cells and AMs. Tumor cell secreted GM-CSF leads to the proliferation and accumulation of metabolically and immunologically supportive AMs. GM-CSF signaling through PPARγ facilitates AM adaptation to the TME by increasing lipid metabolism and inhibiting inflammation. In turn, AMs increase cholesterol efflux making cholesterol available to tumor cells, which then utilize the nutrient to support optimal EGFR-oncogenic signaling. This study identifies a novel paradigm by which tumor cells co-opt AM homeostatic cytokine and lipid signaling pathways to establish an immunologically and metabolically permissive TME. Furthermore, it identifies PPARγ antagonists and statins as promising immuno-metabolic targeted therapies for revitalizing T cell effector functions in the lungs while also exploiting vulnerabilities in oncogenic EGFR signaling.

## Results

### EGFR-mutant LUADs contain an abundance of tissue-resident alveolar macrophages due to increased GM-CSF signaling from tumor epithelium

Although EGFR^+^ lung cancer tumors are susceptible to EGFR targeted kinase inhibitor (TKIs), they present with low tumor mutation burden, are devoid of T cells, and are unresponsive to anti-PD1/PD-L1 immune checkpoint blockade (ICB) (5). To devise tailored immunotherapeutic strategies and to better understand how the immune landscape evolves throughout EGFR^+^ LUAD tumor development, we profiled the immune infiltrate using flow cytometry across different phases of tumor progression (**Fig S1A**) in a previously described *Ccsp-rtTA;TetO-EGFR^L858R^* bi-transgenic mouse model of EGFR-driven LUAD (**Fig 1A–B**) that recapitulates critical features of human disease (40). Consistent with our previous report (39), we observed a decrease in the density of CD8^+^ and CD4^+^ T cells as tumors progressed (**Fig 1C**). In contrast, there was a dramatic increase in the lung-resident SiglecF^+^ CD11c^+^ CD11b^-^ alveolar macrophage (AM) compartment (39,40). After 4-5 weeks of tumor growth, AMs comprised ∼70-75% of the CD45^+^ cells in the lungs, whereas other myeloid populations such as CD11b^+^F4/80^+^ interstitial macrophages (IMs) and CD11b^+^ Ly6C^mid^ and Ly6C^hi^ monocytes remained unchanged or decreased as tumors progressed (**Fig 1C and S1B**). This demonstrates that tissue-resident AMs expand concurrently with transformed AT2 ‘client’ cells, suggesting a tight homeostatic relationship between the two cell types.

**Figure 1.**
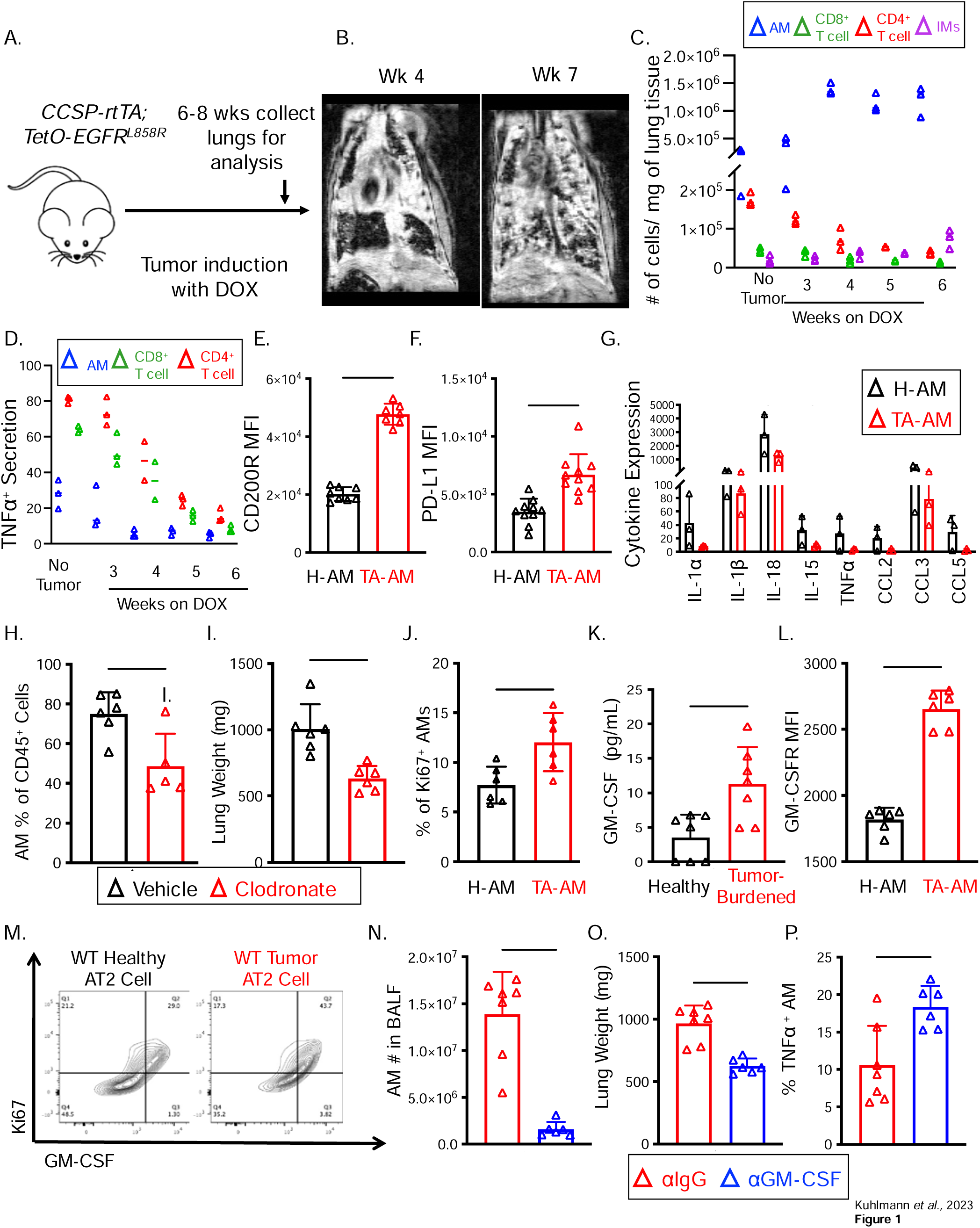
Alveolar Macrophages accumulate during lung tumorigenesis, become increasingly tolerogenic, and promote LUAD growth. Schematic of lung adenocarcinoma (LUAD) induction and immune profiling in a genetically inducible EGFR^L858R^ mouse LUAD model. Tumors were initiated via feeding LUAD mice (*Ccsp-rtTA; TetO-EGFR^L858R^*) or littermate control ‘Healthy’ mice (*TetO-EGFR^L858R^* or *Ccsp-rtTA*) doxycycline (DOX) in chow diet. Unless otherwise noted, mice were analyzed 6-8 weeks on DOX. (B) Serial MRI of a mouse after 4 and 6 or 7 weeks on DOX. (C) Immune infiltrates were quantified by flow cytometry during disease initiation (2-3 weeks), at emergence of macroscopic disease (4 weeks) and with fully established disease (6-8 weeks). Alveolar macrophages (AMs) were defined as CD45^+^CD11b^-^ SigF^+^CD11c^+^and interstitial macrophages (IMs) were defined as CD45^+^AM^-^CD11b^+^Ly6G^-^. CD4^+^ and CD8^+^ T cells were gated on CD45^+^CD3^+^ cells. (D) as in C, but AMs were stimulated with LPS and T cells were stimulated with PMA and ionomycin for 6 hrs prior to intracellular TNF staining. (E-F) After 6-8 weeks on DOX, AMs were isolated from LUAD mice (red) or littermate controls (black) and amounts of the inhibitory receptor CD200R (E) or checkpoint ligand PD-L1 (F) were measured by flow cytometry based on mean fluorescent intensity (MFI). (G) Bulk RNA-sequencing of AMs from late stage EGFR^L858R^ lung tumors were analyzed for expression of several pro-inflammatory cytokines (n=3). Data from Ayeni et al. 2019. (H-I) LUAD mice were administered clodronate liposomes 2× weekly retroorbitally at weeks 4-6 of DOX and then sacrificed at week 6 of Dox. AM frequency was measured by flow cytometry (H) or tumor burden was assessed by the dry lung weight (I). (J) AM proliferation rates were assessed by Ki67-staining and flow cytometry. (K) GM-CSF was measured in healthy and LUAD lung lysates using ELISA. (L) AM surface expression of GM-CSFR was measured using flow cytometry. (M) Representative flow plots showing intracellular GM-CSF and Ki67 in healthy and malignant AT2 cells (CD45^-^Epcam^+^CD31^-^MHCII^hi^proSPC^+^) incubated with BFA for 6 hours. (N-P) GM-CSF blocking (blue) or isotype control mAbs (red) were administered twice weekly (0.5mg/mouse) i.p. to LUAD mice during weeks 4-7 on DOX. We then measured the number of TA-AMs present in the BALF fluid (N), tumor burden using lung dry weight as a surrogate (O), and AM TNF secretion following LPS stimulation (P). Data shown are mean ± SEM, and statistical analysis were performed by two-tailed unpaired Student’s test (E-P). *p<0.05, **p < 0.01, ***p<0.001, ****p < 0.0001. Data are representative from ≥3 experiments (C,D,M) or pooled from ≥3 experiments (E-F,H-L,N-P) with each group containing 3 (C,D), 8-7 (E), 10 (F), 6-5 (H), 6 (I-J,L), 7 (K), or 7-6 (N-P) mice.

Given the profound LUAD-driven expansion of tumor associated AMs (TA-AMs) we next interrogated their functional properties. Compared to AMs found in healthy lungs (H-AMs), the TA-AMs were skewed towards a more tolerogenic state based on *(i)* lower TNF production following LPS stimulation (**Fig 1D, S1C**), *(ii)* higher surface expression of the inhibitory receptor CD200R (**Fig 1E, S1E**) and the checkpoint molecule PD-L1 (**Fig 1F, S1G),** and *(iii)* diminished mRNA expression of several pro-inflammatory cytokines and T cell chemo-attractant chemokines (**Fig 1G**). The few remaining T cells also displayed reduced effector cytokine secretion (**Fig 1D**), but the IMs and monocytes were not as visibly affected functionally (**Fig S1D-H**).

To assess if the accumulation of tolerogenic TA-AMs was incidental or necessary for tumor growth, we partially depleted the cells using clodronate liposomes. Compared to vehicle-treated tumor-bearing lungs, clodronate liposome treatment significantly decreased the number of TA-AMs (**Fig 1H**) and this corresponded with a significant decrease in tumor burden (**Fig 1I**) in agreement with previous reports, indicating that AMs promote tumorigenesis (33,40). These results suggest that AMs-but not other myeloid cells-expand and possess tolerogenic properties as LUAD progresses and targeting their accumulation represents an immunotherapeutic target for EGFR-driven LUAD.

GM-CSF and IL-4 were both candidates for AM accumulation in the TME because tonic epithelial AT2-derived GM-CSF is required for AM development/maintenance (41) and both cytokines induce AM proliferation (42,43). Indeed, the increase in Ki-67^+^ TA-AMs correlated with greater amounts of GM-CSF, but not IL-4 in tumor lysates (**Fig 1J–K** and data not shown). Accordingly, TA-AMs had increased surface expression of the GM-CSF receptor CD116 relative to H-AMs (**Fig 1L**). In homeostasis, AMs respond exclusively to GM-CSF from AT2 cells, despite production from hematopoietic sources (41). Consistent with previous reports (40) AT2 cells from tumor bearing lungs increased their GM-CSF secretion (**Fig 1M**). To more directly test if GM-CSF was required for AM accumulation in the TME, GM-CSF was blocked by administration of αGM-CSF monoclonal antibody (mAb) or isotype control mAbs intra-peritoneal (I.P.) from weeks 3-6 after tumor initiation when the burst of TA-AMs is most evident. The αGM-CSF treatment caused a dramatic reduction of TA-AMs in the bronchial alveolar lavage fluid (BALF) and to a lesser extent in the lungs relative to controls (**Fig 1N, S1I**) and significantly reduced lung weight (**Fig 1O**). TA-AMs from GM-CSF depleted lungs were also more pro-inflammatory than their isotype control treated counterparts, with a greater capacity to secrete TNF following LPS stimulation (**Fig 1P, S1J**). In contrast, we did not see appreciable differences in either infiltration or inflammatory cytokine secretion from IMs or monocytes following GM-CSF depletion (**Fig S1I-K**). These data show that AMs expand and promote LUAD growth in a GM-CSF-dependent fashion.

### Metabolic reprogramming of AMs in LUAD lungs containing increased lipids and surfactant

Surfactant homeostasis in the lungs is maintained by a delicate balance of catabolism and recycling undertaken by both AT2 cells and AMs. Lung surfactant lipids and proteins are a defining feature of the alveolar niche with known immunoregulatory roles however, it is unknown how transformation of surfactant producing AT2 cells might disrupt this balance and therefore we set out to define the alterations that transformation has on the lung surfactant system. To examine if surfactant proteins and lipids were altered in patients with lung cancer (**Table S1**) we compared the amounts of Surfactant Protein D (SP-D) and lipids in the bronchial alveolar lavage fluid (BALF) of patients diagnosed with Lung Cancer to those diagnosed with other inflammatory lung diseases such as COPD or a mycobacterial infection (**Fig 2A, E**). A similar analysis of BALF from EGFR^+^ LUAD-bearing mice was also performed (**Fig 2B, F**). This revealed significant increases in hSPD or mSPA and mSPD in both human (as compared to infection) and murine LUAD specimens, respectively. Further, mRNA analysis of epithelium from murine EGFR^+^ LUAD showed that all four SP genes (*S*ftpa1*, b, c, d)* had elevated expression compared to lungs of healthy littermates (**Fig 2C**). To examine if surfactant protein expression was incidental or helped contribute to tumorigenesis, we generated a strain of EGFR^+^ LUAD mice with deletion of surfactant proteins A and D (SftpAD-/-tumor mice) and observed a significant decrease in lung weight as compared to littermate WT controls (**Fig 2D**). Lipidomic analysis demonstrated highly consistent increases in surfactant lipids, such as Free Fatty Acids (FFAs), Cholesterol Esters (ChEs) and phospholipids, in BALF, in both human and mouse tumor-bearing lungs relative to those with an infection or COPD (human) or healthy lungs (mouse) (**Fig 2E–F**). These results point out that increased surfactant proteins and associated lipids are features of lung cancer and likely their accumulation promotes LUAD growth.

**Figure 2.**
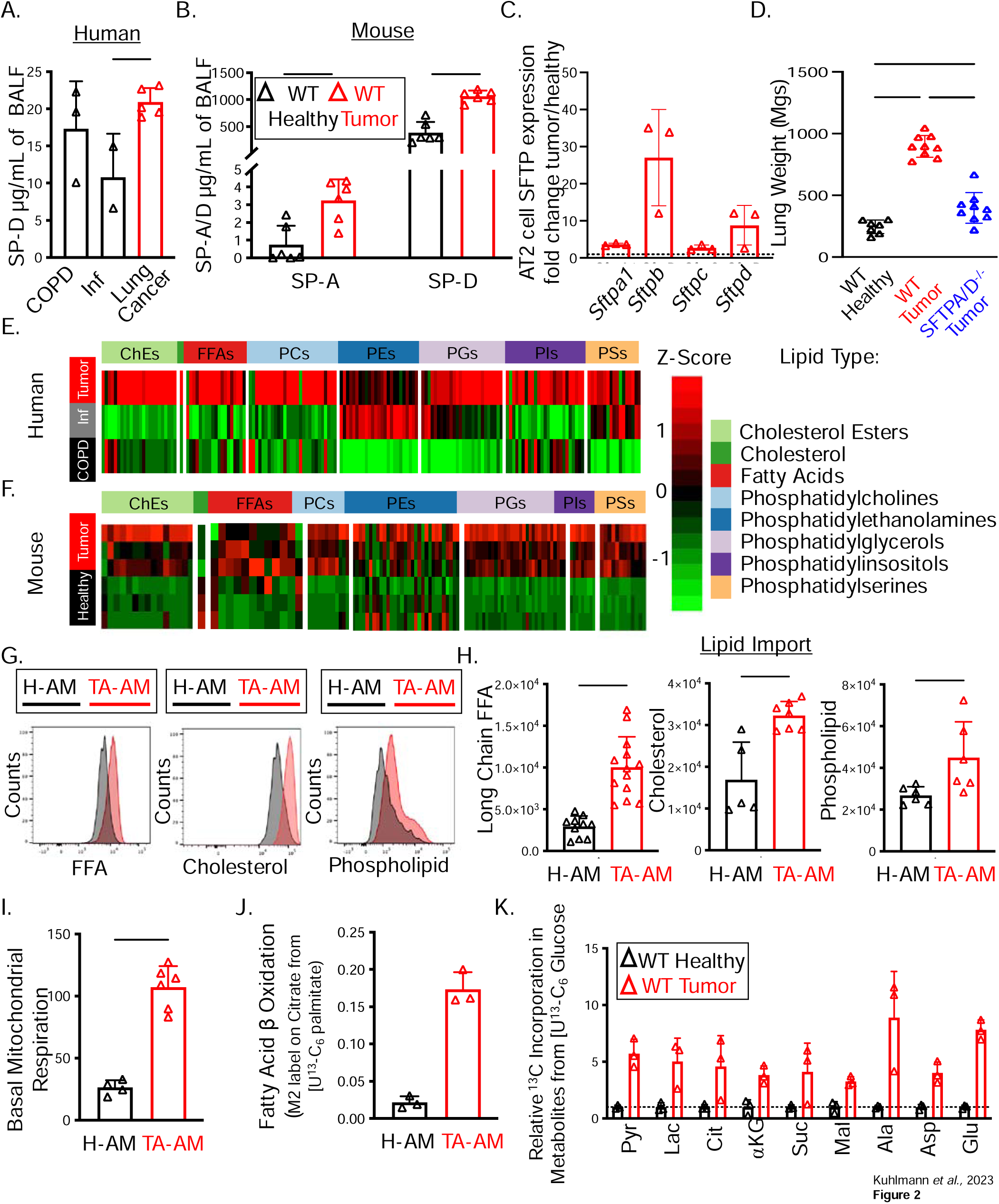
Lung surfactants accumulate in the TME despite increased levels of AM lipid uptake. (A) Bar graphs show SP-D concentrations in BALF collected from patients undergoing diagnostic lung lavages for lung cancer (n=5), COPD (n=3), or mycobacterial infection (n=2) as measured by ELISA. (B) Concentration of SP-D and SP-A from the BALF of littermate control (WT Heathy, black) mice or those with LUAD (WT Tumor, red) as measured by ELISA. (C) Bar graph shows the fold change in surfactant proteins mRNAs from EGFR^L858R^ lung epithelium (EPCAM^+^) in LUAD lungs (39). (D) SPA^-/-^ SPD^-/-^ double knockout mice were crossed with *Ccsp-rtTA;TetO-EGFR^L858R^* mice and placed on DOX for 6-8 weeks and dry lung weight was measured. (E-F) Similar to (A-B), heatmaps show lipids in the BALF from patients with lung cancer (red), COPD (black), or mycobacterial infection (gray) (E) or from littermate control (WT Heathy, black) mice or those with LUAD (WT Tumor, red) (F) as measured by LC/MS. Heatmaps depict the relative abundance of each class of lipid species normalized to volume (shown as a row Z score). Human data depict abundance averaged amongst the samples. (G-H) Import of free fatty acids, cholesterol and phospholipids (Bodipy C12, NBD cholesterol, DPPE respectively) were compared between AMs isolated littermate controls (H-AM, black) and LUAD mice (TA-AM, red) using flow cytometry and displayed as representative histograms of lipid import (G) and cumulative bar graphs of MFI (H). (I-K) H-AMs and TA-AMs were isolated 6-8 weeks on DOX and rates of basal mitochondrial respiration was measured using Seahorse Flux Analyzer or cells were labeled with ^13^C-palmitate (J) or ^13^C-glucose (K) and measured for rates of fatty acid oxidation and other metabolites, respectively. Data shown are mean ± SEM, and statistical analysis were performed with a two-tailed unpaired Students test (A-C, H-K) or a two way ANOVA (D). *p<0.05, **p < 0.01, ***p<0.001, ****p < 0.0001. Data are pooled from ≥3 experiments with each group containing 6 (B), 7-9 (D), 5-13 (H) animals or is representative from ≥2 experiments with each group containing 3 (G,I-K) animals. Human samples were collected from 3 (COPD), 2 (mycobacterial infection) or 5 (lung cancer) and data was pooled (A) or averaged (D).

Given that surfactant associated lipids were elevated in tumor BALF and that AMs play a critical role in regulating lipid levels in the lungs, we next investigated how the TA-AMs adapted to this lipid-rich TME. Conceivably, lipid accumulation in the lungs could occur if TA-AMs stopped or reduced lipid import, however, flow cytometric measurements of import of fluorescent lipid analogues showed the opposite. That is, TA-AMs bound to and imported more long chain FFAs, cholesterols, and phospholipids than their H-AM counterparts (**Fig 2G–H**). Significant increases in lipid import were most apparent in AMs as neither Ly6C^mid^ nor Ly6C^hi^ monocytes displayed increased import of lipids and IMs only increased import of cholesterol but not FFA or phospholipids (**Fig S2**). Furthermore, compared to the H-AMs, TA-AMs displayed an increase in lipid and mitochondrial flux based on *(i)* heightened mitochondrial respiration by seahorse (**Fig 2I),** *(ii)* increased fatty acid oxidation using ^13^C labelled palmitate (**Fig 2J**), *(iii)* and an increase in ^13^C-glucose labelling on TCA cycle intermediates (**Fig 2K**). Altogether, these data demonstrate that TA-AMs were not only the most supernumerary cell type, but were also metabolically reprogrammed in LUAD through increased rates of surfactant lipid import and breakdown, glycolysis and mitochondrial flux.

### Inhibition of PPAR**γ** in TA-AMs suppresses LUAD growth

PPARγ is a central regulator of lipid metabolism and alternative activation in macrophages (44,45). Specifically in AMs, PPARγ acts downstream of GM-CSF signaling to govern AM differentiation (46). Humans whose AMs do not upregulate PPARγ due to autoantibodies against GM-CSF develop alveolar proteinosis (PAP), a disease marked by disrupted surfactant homeostasis and steady state lung inflammation. As such, we postulated that PPARγ may be involved in the changes in TA-AMs that promoted LUAD growth. In support of this hypothesis, previously published CyTOF on NSCLC samples reported an increase in PPARγ^+^ macrophages infiltrating patient tumors relative to adjacent normal tissue but did not specify in which macrophage subset PPARγ^+^was acting (47). Therefore, we reanalyzed previously published single cell RNA sequencing (scRNA-seq) data (48) and CyTOF data (47) from LUAD tumors and matched adjacent healthy tissue and confirmed that AMs were the highest expressors of PPARγ in myeloid cell subsets and that expression of PPARγ increased during tumorigenesis in AMs and IMs but not in monocytes (**Fig 3A–B, S4B**).

**Figure 3:**
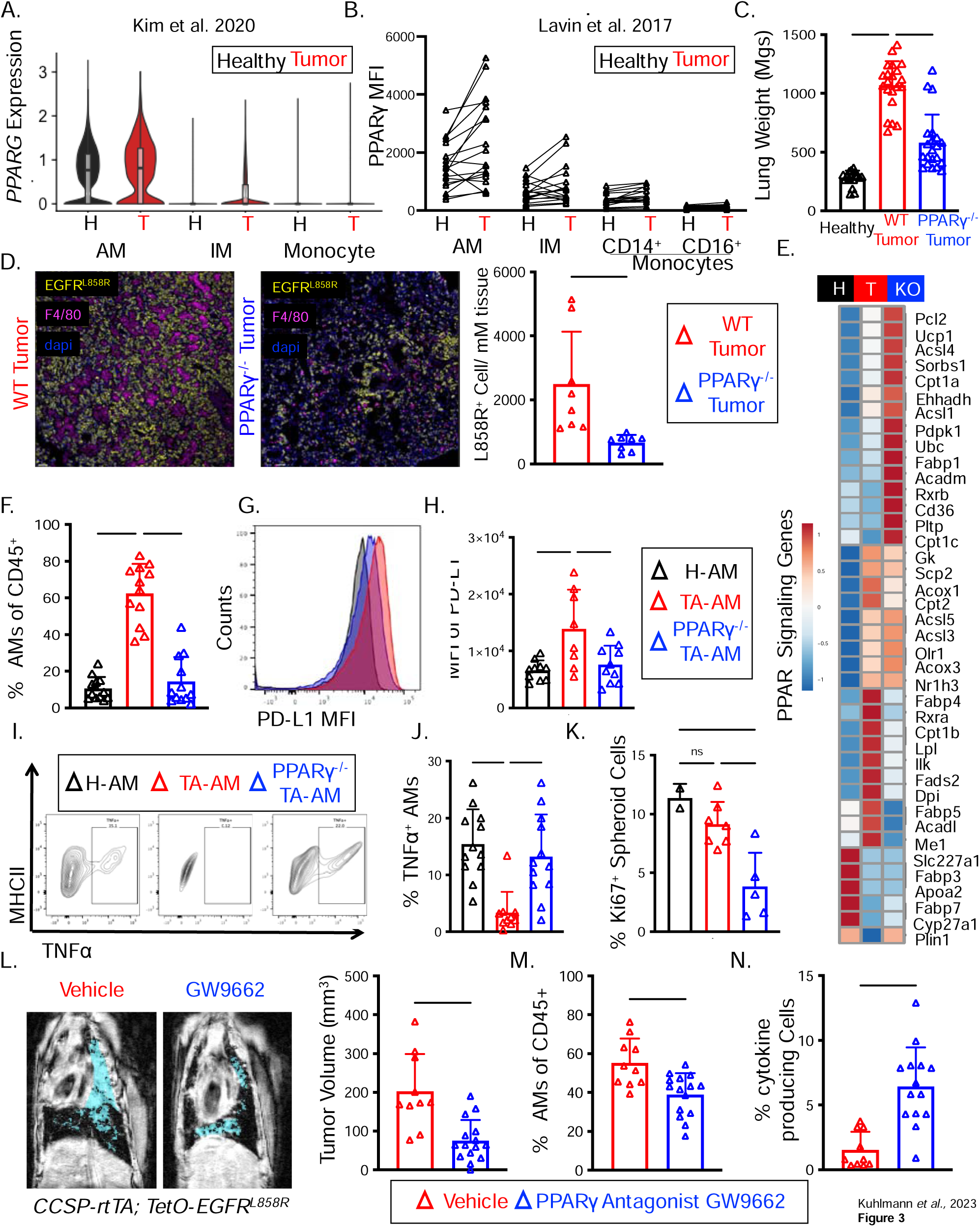
AM PPARγ is required for the accumulation and phenotype in the TME. (A) Violin plots of of *PPARG* mRNA in macrophage subsets from patients with NSCLC or matched healthy adjacent tissue (data from (48) GSE131907). (B) MFI of PPARγ in various myeloid subsets in paired samples from NSCLC tumors or healthy adjacent tissue from the same patient measured by CYTOF analysis (data from Lavin et al. 2017). The following gating strategy was used: AMs: CD11b^+^CD64^+^CD163^+^CD206^hi^, IMs: CD11b^+^CD64^+^CD163^-^, CD14^+^ Monocytes: CD11b^+^CD64^-^CD14^+^, and CD16^+^ Monocytes: CD11b^+^CD64^-^CD16^+^. (C-G) LUAD mice with deletion of PPARγ in macrophages (*Pparγ^Fl/Fl^; Csf1r^Cre^, Ccsp-rtTA; TetO-EGFR^L858R^)* (referred to as PPARγ Tumors, blue) and littermate controls (*Ppar*γ*^Fl/F^; Ccsp-rtTA; TetO-EGFR^L858R^)* (referred to as WT Tumors, red) were placed on DOX for 7-9 weeks at which point tumor burden and immune infiltrates were examined by scRNA-seq, microscopy and flow cytometry. Healthy lungs (black) were from control *TetO-EGFR^L858R^* mice on DOX. (C) Bar graph of dry lung weights. (D) Immunofluorescence microscopy was performed on lung sections from WT Tumor (red) and PPARγ^-/-^ Tumor (blue) measuring density of tumor cells (EGFR^L858R^), AMs (F4/80) and nuclei (dapi). (E) Heatmap shows Z-score by row of PPAR-target gene expression (from scRNA-seq) in the AM cluster from healthy (H, black) lungs or those with LUAD from WT (T, red) and PPARγ^-/-^ (KO, blue) Tumors. (F-H) Percentage of AMs (F) and MFI of PD-L1 on AMs (G,H) as assessed by flow cytometry from the three groups of mice. (I-J) H-AMs (black) or TA-AMs from WT (red) and PPARγ^-/-^ (blue) Tumors were isolated and stimulated with LPS to measure TNF production by flow cytometry. (K) Spheroids from human EGFR^del19^ cell line HCC827 were cultured on low attachment plates for a week and then AMs isolated from healthy lungs or those with LUAD from WT (red) or PPARγ^-/-^ (blue) Tumors were added with GM-CSF (20 pg/mL) for three days and proliferation was measured by ki67 staining. (L-N) LUAD mice and littermate controls were placed on DOX for three weeks and then treated by oral gauvage 5x/week with the PPARγ antagonist GW9662 (1mg/kg) in corn oil or vehicle alone for three more weeks. (L) Representative MRI images and quantification of tumor burden in untreated and antagonist treated lungs. Percentage of AMs (M) and those producing TNF after LPS stimulation (N) were assessed by flow cytometry. Data shown are mean ± SEM, and statistical analysis were performed with a two-tailed unpaired Students test (D,L-N) or a two way ANOVA (C,F,H,J-K). *p<0.05, **p < 0.01, ***p<0.001, ****p < 0.0001. Data are pooled from ≥3 experiments with each group containing 13 (B) patients and 4 (D), 17-21 (F), 12 (H), 8-10 (J) 10-12 (K), 10-15 (L,M) and 10-14 (N) mice.

To examine the role of PPARγ in TA-AM metabolic reprogramming, we aimed to delete PPARγ from AMs, however, there is currently no AM-specific Cre transgenic murine strain available. Therefore, we generated two different strains of EGFR^+^ LUAD mice that enabled conditional deletion of PPARγ in different myeloid compartments: a *Csf1r*^Cre^ for deletion in macrophages and a *Cd11c*^Cre^ for deletion in CD11c^+^ AMs (46) as well as other conventional dendritic cells (cDCs). Previous reports showed that deleting PPARγ using either cell types significantly impaired development of AMs, but did not affect IM, monocyte, or cDC accumulation or phenotype in the lungs (46). Therefore, we closely compared both models (*Ccsp-rtTA;TetO-EGFR^L858R^*; *Ppar*γ*^fl/fl^;Csf1r^Cre^*and *Ccsp-rtTA;TetO-EGFR^L858R^*; *Ppar*γ*^fl/fl^;Cd11c^Cre^*) to identify reproducible TA-AM and myeloid phenotypes in LUAD. We refer to the TA-AMs analyzed from the tumors in these mice as ‘**PPAR**γ**^-/-^ TA-AMs**’ and distinguish them from the *wild type* ‘**WT TA-AMs’** in littermates lacking a Cre transgene but still expressing *Ccsp-rtTA;TetO-EGFR^L858R^* as well as the *healthy* **H-AMs** found in littermates lacking both Cre and *EGFR^L858R^* transgenes. Total tumor bearing lung samples from these mice are respectively referenced as ‘**PPAR**γ**^-/-^ Tumor**’, ‘**WT Tumor**’ and ‘**WT Healthy**’.

First and foremost, in both models, after 8-10 weeks of DOX administration we observed reduced lung weights (**Fig 3C and S3A**) and density of EGFR^L858R+^ cells in PPARγ^-/-^ Tumors (**Fig 3D**) compared to WT Tumors, indicating that PPARγ is required in TA-AMs and possibly other myeloid cells to support tumor growth. Second, single cell RNA-sequencing (scRNA-seq) on cells from the *Ccsp-rtTA;TetO-EGFR^L858R^*; *Ppar*γ*^fl/fl^;Csf1r^Cre^* mice confirmed that while PPARγ^-/-^ TA-AMs still clustered with WT TA-AMs (**Fig S4A**), they had decreased expression of several PPAR target genes (**Fig 3E**). Third, as expected, the lungs of mice that lacked PPARγ in AMs contained very few mature SiglecF^+^ CD11c^+^ CD11b^-^ AMs, but had a larger number of SiglecF^+^ CD11c^+^ CD11b^+^ immature AM-like (im-AM) cells (**Fig 3F and S3B**) and a subtler increase in interstitial macrophage and monocytes (**Fig S3B**). Fourth, there were also increased numbers of tumor infiltrating CD8^+^ and CD4^+^ T cells in PPARγ^-/-^ Tumors *vs.* WT Tumors (**Fig S3C**), indicating the tumors were more inflamed in mice that lacked PPARγ in the AMs or macrophages. Fifth, flow cytometry analysis revealed that PPARγ^-/-^ TA-AMs produced lower amounts of the inhibitory ligand PD-L1, but higher amounts of TNF than the WT TA-AMs (**Fig 3G–J**), and this effect was fairly selective to the TA-AMs as these properties were not significantly altered in the IMs, monocytes, DCs or T cells (**Fig S3D-F**). These findings demonstrate that PPARγ is necessary for sustaining the increased numbers of anti-inflammatory, tolerogenic state of TA-AMs in LUAD. Lastly, using 3-D culture models, we compared the ability of murine WT TA-AMs and PPARγ^-/-^ TA-AMs to support the growth of human EGFR^Del19^ tumor spheroids (cell line HCC827) and found that WT TA-AMs supported spheroid proliferation and growth significantly better than PPARγ^-/-^ TA-AMs (**Fig 3K**). Collectively, these data show that mature TA-AMs support LUAD growth and that increased PPARγ activity in these cells contributes to their tumor supportive and anti-inflammatory roles.

Both methods of genetic deletion of PPARγ in TA-AMs profoundly suppressed tumor growth, but a potential caveat is that these genetic systems also interfere with AM development prior to and throughout the course of tumor progression. Therefore, we next wanted to investigate if there was any therapeutic effect to inhibiting PPARγ after tumor initiation in animals that contain mature TA-AMs. To this end, we treated EGFR^+^ LUAD mice on DOX with the PPARγ antagonist GW-9662 via oral gavage from weeks 3-6 and analyzed the mice at 6 weeks. Magnetic Resonance Imaging (MRI) revealed that GW-9662 treatment significantly lowered tumor burden (**Fig 3L**) and TA-AM frequencies (**Fig 3M**), yet increased TNF secretion in TA-AMs (**Fig 3N**). Consistent with what we observed in the two genetic models above, inhibition of PPARγ by GW-9662 did not alter IM or monocyte recruitment or their inflammatory states (**Fig S3G-H**). These promising results suggest that PPARγ inhibition could be a worthy immunotherapeutic target for limiting EGFR^+^ tumor growth while reversing TA-AM tumor-promoting states.

### PPAR**γ** metabolically reprograms TA-AMs and controls their tumor-promoting properties

In addition to controlling macrophage inflammatory states, PPARγ is a master regulator of macrophage lipid metabolism. In patients with PAP, impaired PPARγ expression in AMs hinders their surfactant catabolism resulting in AM foam cell formation, deregulated lung metabolism and steady-state lung inflammation (49). Given that loss of PPARγ in TA-AMs slowed tumor growth we reasoned that perhaps PPARγ directs the metabolic alterations in TA-AMs, fueling their tumor supportive properties. We first compared the lipidomes of H-AMs, WT TA-AM, and PPARγ^-/-^ TA-AMs using mass spectrometry and this showed that PPARγ^-/-^ TA-AMs become vastly lipid laden, accumulating both neutral lipids and phospholipids relative to AMs found in LUAD or healthy lungs (**Fig 4A**). Further, GSEA of scRNA-seq data in mice and humans identified that the expression of several genes involved in fatty acid and cholesterol metabolism and oxidative phosphorylation were increased in WT TA-AMs and human TA-AMs from NSCLC (**Fig 4B–C** and **S4C-E**) **(48**) relative to healthy AMs, and this was dependent on PPARγ in the murine TA-AMs. To examine the metabolic activity of TA-AMs directly, we measured rates of fatty acid import, synthesis and oxidation in TA-AMs using C12-Bodipy,^14^C acetate and ^14^C palmitate, respectively. This showed that unlike the WT TA-AMs, the PPARγ^-/-^ TA-AMs did not display increased FFA uptake, synthesis or oxidation (**Fig 4D–E,G**). Further, while WT TA-AMs had significantly elevated mitochondrial respiration and TCA cycle flux (based on ^13^C glucose tracing), this was not observed in the PPARγ^-/-^ TA-AMs (**Fig 4H–I**). These results demonstrate that PPARγ is a central regulator of the metabolic rewiring of TA-AMs in LUAD and that many metabolic changes were normalized by loss of PPARγ, correlating with slower tumor growth.

**Figure 4:**
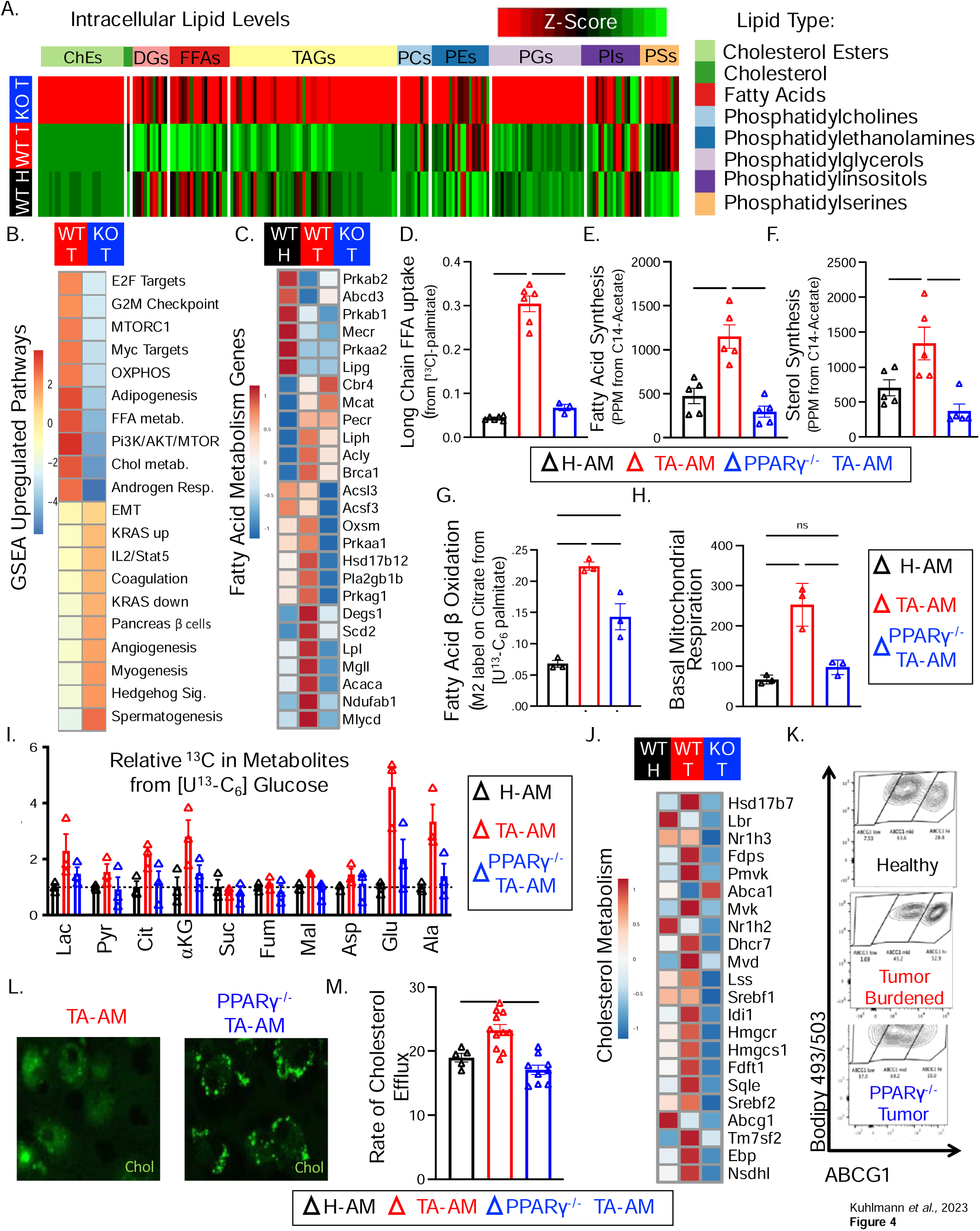
PPARγ rewires Alveolar macrophage metabolism within the TME. LUAD mice lacking PPARγ in macrophages (PPARγ^-/-^ Tumors or KO Tumors (blue)) and littermate ‘WT Tumor’ controls (*Ppar*γ*^Fl/F^; Ccsp-rtTA; TetO-EGFR^L858R^*(red)) along with ‘WT Healthy’ lung controls (*TetO-EGFR^L858R^* or *Ccsp-rtTA* (black)) were placed on DOX for 7-9 weeks. (A) Heatmap depicts the relative abundance (averaged across 3 samples/group) of the indicated lipid species in BALF normalized to the total volume as measured by LC/MS. (B) WT and PPARγ^-/-^ KO TA-AMs were compared by scRNA-seq to identify the top 10 differentially upregulated pathways using GSEA. (C) Heatmap shows expression of fatty acid metabolism genes by row Z-score (from scRNA-seq) in WT AMs from Healthy lungs (black) *vs.* WT (red) and PPARγ^-/-^ (KO, blue) TA-AMs from LUAD lungs. (D-I) AMs isolated from WT Healthy lungs (black) and TA-AMs isolated WT (red) or PPARγ^-/-^ (blue) Tumors were incubated with (D) [^13^C]-palmitate, (E) [^14^C]-acetate, (F) [^14^C]-acetate, [^14^C]-palmitate, or (I) [^13^C]-glucose and assayed for (D) free fatty acid (FFA) import, (E) FA synthesis, (F) sterol synthesis, (G) FA b-oxidation (I) or relative flux into several metabolites. (H) Basal mitochondrial respiration in AMs was performed using seahorse extracellular flux assay. (J) Heatmap shows expression of cholesterol synthesis and metabolism genes by row Z-score (from scRNA-seq) in WT AMs from Healthy lungs (black) *vs.* WT (red) and PPARγ^-/-^ (KO, blue) TA-AMs from LUAD lungs. (K) ABCG1 expression on AMs was assessed by flow cytometry. (L-M) Control and PPARγ^-/-^ TA-AMs were cultured with NBD-Cholesterol (green) for one hour and then (L) imaged by confocal microscopy to examine lipid droplets. To assess cholesterol efflux (M), the labeled TA-AMs and H-AMs were then equilibrated overnight in serum free media followed by incubation with FBS as a cholesterol acceptor for 4 hours and effluxed NBD-Cholesterol in the supernatant was measured by fluorescence. Data shown are mean ± SEM, and statistical analysis were performed with a two way ANOVA. *p<0.05, **p < 0.01, ***p<0.001, ****p < 0.0001. Data are representative from 2 experiments with an N=3 (A). Data are pooled from ≥2 experiments with each group containing 6 (D), 5 (E, F), 3 (G-H), 5-6 (I) or 5-11 (M) animals.

Studies on PAP point to a pivotal role for cholesterol in driving global AM metabolic dysfunction in these patients (50). As PPARγ^-/-^ TA-AMs had significantly increased storage of ChEs by mass spectrometry (**Fig 4A),** we aimed to further characterize their cholesterol metabolism. Single cell RNA sequencing revealed that while WT TA-AMs and human TA-AMs upregulated genes involved in cholesterol synthesis and metabolism, this was abrogated in mouse PPARγ^-/-^ TA-AMs (**Figs 4J** and **S4E**). A similar pattern of increased sterol synthesis was confirmed in the WT TA-AMs relative to the H-AMs and PPARγ^-/-^ TA-AMs using ^14^C acetate tracing into sterol lipid fractions (**Fig 4F**). While there was not a difference in cholesterol import in PPARγ^-/-^ TA-AMs as compared to WT TA-AMs (data not shown) there was a decrease in the cholesterol efflux transporter ABCG1 on these cells (**Fig 4J, K**) and when fed with labelled cholesterol, PPARγ^-/-^ TA-AMs robustly stored the esterified cholesterol in lipid droplets (**Fig 4L**). A fluorometric cholesterol efflux assay confirmed that while WT TA-AMs increased their rates of cholesterol efflux relative to **H-AMs**, PPARγ^-/-^ TA-AMs did not (**Fig 4M**). Collectively, these results show that PPARγ acts a rheostat, balancing rates of cholesterol synthesis, esterification and storage *vs.* efflux in TA-AMs and that elevated PPARγ activity in TA-AMs potentiates cholesterol efflux, anti-inflammatory states and tumor growth.

### GM-CSF—PPAR**γ** signaling axis promotes metabolic and functional reprogramming of TA-AMs to regulate tumor cell cholesterol metabolism

Prior reports have shown that GM-CSF can regulate cholesterol efflux and surfactant lipid import in AMs (50), therefore, we next set out to examine if GM-CSF also regulates the phenotypic and metabolic rewiring of TA-AMs in a PPARγ-dependent manner. We cultured purified WT TA-AMs and PPARγ^-/-^ TA-AMs overnight in the presence or absence of a physiological dose of GM-CSF (20 pg/mL, **Fig 1M**) and observed that GM-CSF induced PD-L1 upregulation (**Fig. 5A**). Metabolically, GM-CSF increased fatty acid synthesis and cholesterol efflux, yet decreased cholesterol esterification in WT TA-AMs, however, this was not observed in PPARγ^-/-^ TA-AMs (**Fig 5B–D**). Notably, the PPARγ^-/-^ TA-AMs had substantially increased rates of cholesterol esterification that correlated with increased ChE storage (**Figs. 5D and 4A, K**). Given that blocking GM-CSF suppressed tumor growth similar to deleting PPARγ in TA-AMs (compare **Figs 1O and 3C, S3A**), we reasoned that GM-CSF-mediated cholesterol efflux from TA-AMs may be aiding tumor growth by providing cholesterol to meet the heightened metabolic demands of rapidly proliferating tumor cells. To directly query if cholesterol effluxed from TA-AMs could be taken up by AT2 cells, we first pulsed TA-AMs with Bodipy-cholesterol and then co-cultured them with EGFR^L858R^ AT2 cells from LUAD (**Fig 5E**). This showed that WT TA-AMs more efficiently transferred labelled-cholesterol directly to tumor cells than PPARγ^-/-^ TA-AMs (**Fig 5F**). These data demonstrate that GM-CSF signals through PPARγ to suppress cholesterol esterification and promote efflux in TA-AMs that can be delivered to tumor cells.

**Figure 5:**
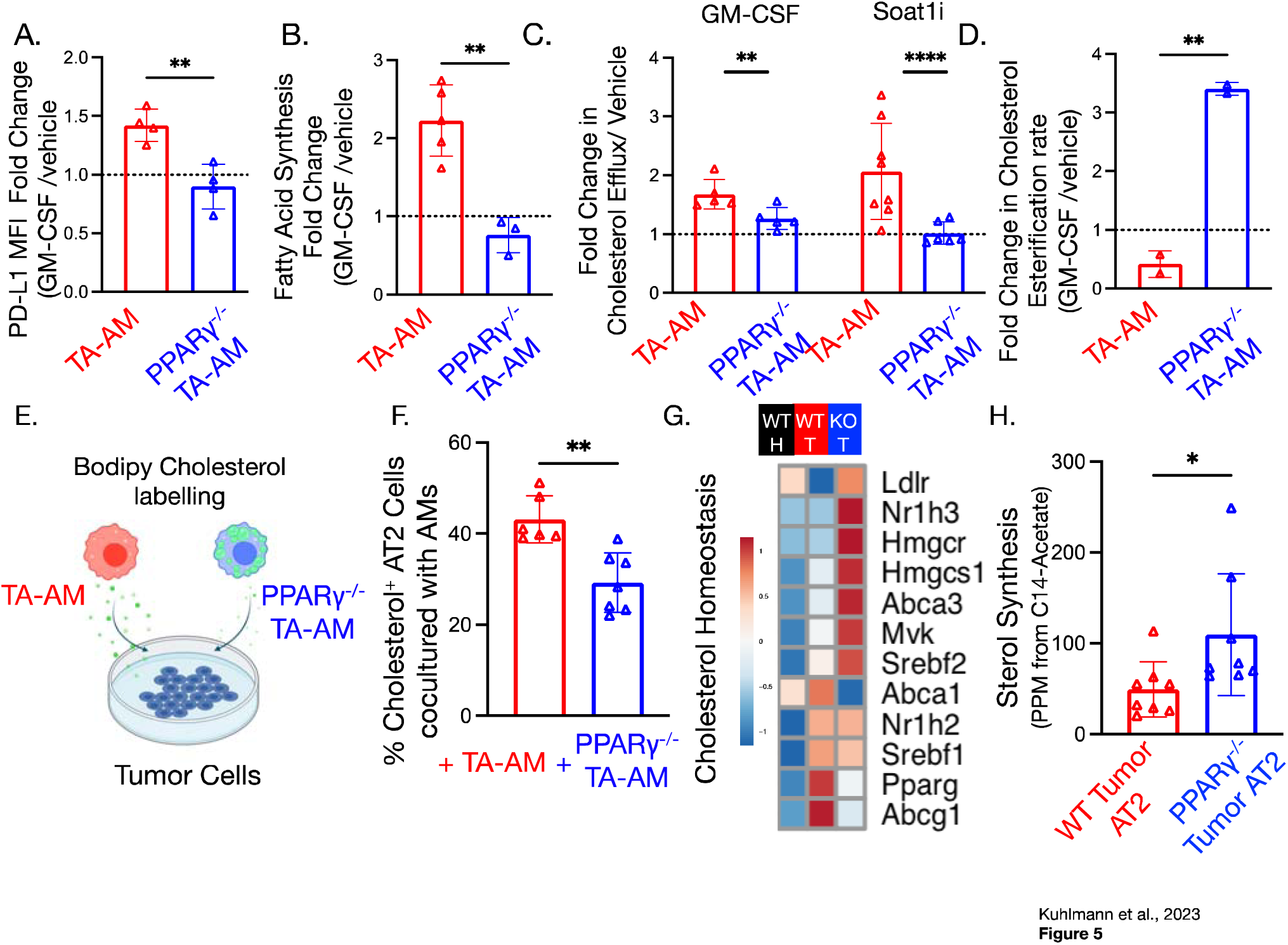
GM-CSF PPARγ signaling drives cholesterol efflux from TA-AMs to tumor cells to promote tumor growth. (A-D) WT and PPARγ TA-AMs were stimulated overnight in the presence or absence of GM-CSF (20 pg/mL) or the SOAT1 inhibitor (Sandoz 58-035 10ug/mL) and assessed for PD-L1 expression by flow cytometry. (A), rates of fatty acid synthesis (B), cholesterol efflux (C), or cholesterol esterification (D). Data were normalized to untreated samples with a dashed line at 1. In (C), cells were also treated with a cholesterol esterification inhibitor SOAT1 to increase efflux. (E-F) Diagram depicting the experimental schema of TA-AM cholesterol transfer experiments wherein TA-AMs were loaded with fluorescent NBD-cholesterol, washed and then co-cultured with dissociated tumor cells overnight (E). Transfer of fluorescently labelled cholesterol from TA-AMs to AT2 cells was assessed by flow cytometry and percentage of AT2 cells with imported NBD-cholesterol is shown in bar graph (F). (G) Heatmap shows expression of cholesterol synthesis and metabolism genes by row Z-score (from scRNA-seq) in AT2 cells isolated from healthy lungs (black) or LUAD lungs that contain WT (red) or PPARγ TA-AMs (blue). (H) Rates of sterol synthesis in sorted AT2 cells isolated from LUAD lungs that contain WT (red) or PPARγ TA-AMs (blue). Data shown are mean ± SEM, and statistical analysis were performed with a two-tailed unpaired Students test. *p<0.05, **p < 0.01, ***p<0.001, ****p < 0.0001. Data are pooled

The ability of TA-AMs to efflux and transfer cholesterol to tumor cells begs the question of whether tumor cells are dependent upon this nutrient support for their growth and what are the effects when this is disrupted by inhibiting PPARγ in TA-AMs? To examine this question we analyzed the changes in AT2 cell gene expression between WT Tumors and PPARγ^-/-^ Tumors via scRNA-seq and found a striking upregulation of cholesterol synthesis gene expression in AT2 cells from PPARγ^-/-^Tumors as compared to WT Tumors (**Fig 5G**). This was confirmed by a heightened incorporation of ^14^C acetate into sterol intermediates demonstrating increased rates of cholesterol synthesis in AT2 tumor cells when the TA-AMs were lacking PPARγ (**Fig 5H**). These data suggest that when TA-AMs are unable to boost cholesterol efflux due to defective PPARγ activity, tumor cells compensate by increasing cholesterol synthesis to obtain necessary sterol intermediates needed for growth and proliferation. This represents an example of metabolic cooperation between tumor cells and macrophages driven by cooption of the GM-CSF—PPARγ—cholesterol efflux axis that turns homeostatic tissue-resident macrophage functions into tumor supportive functions and exposes a novel and targetable axis with therapeutic potential.

### AM PPAR**γ** rewires AT2 metabolism to support oncogenic signaling

We reasoned that the metabolic strain observed in AT2 tumor cells within PPARγ^-/-^ Tumors may affect oncogenic EGFR^L858R^ activity because EGFR signaling directly influences tumor growth and is sensitive to fluctuations in cellular cholesterol metabolism (51). First, we queried our scRNA-seq data, which showed that the expression of several EGFR-signaling target genes (52) upregulated in EGFR^L858R^ AT2 cells in WT Tumors was lower than those in PPARγ^-/-^ Tumors (**Fig 6A**). Reduced EGFR^L858R^ activity was validated by fluorescence microscopy which showed that the intensity of pEGFR staining in proSPC^+^ AT2 cells was decreased in PPARγ^-/-^ Tumor bearing lungs as compared to WT Tumor bearing lungs or healthy lungs (**Fig 6B–C**). These findings reveal that optimal oncogenic EGFR^L858R^ signaling *in vivo* depended on PPARγ-mediated rewiring of TA-AMs, further emphasizing the tight cooperation between TA-AMs and their transformed ‘client’ cells.

**Fig 6:**
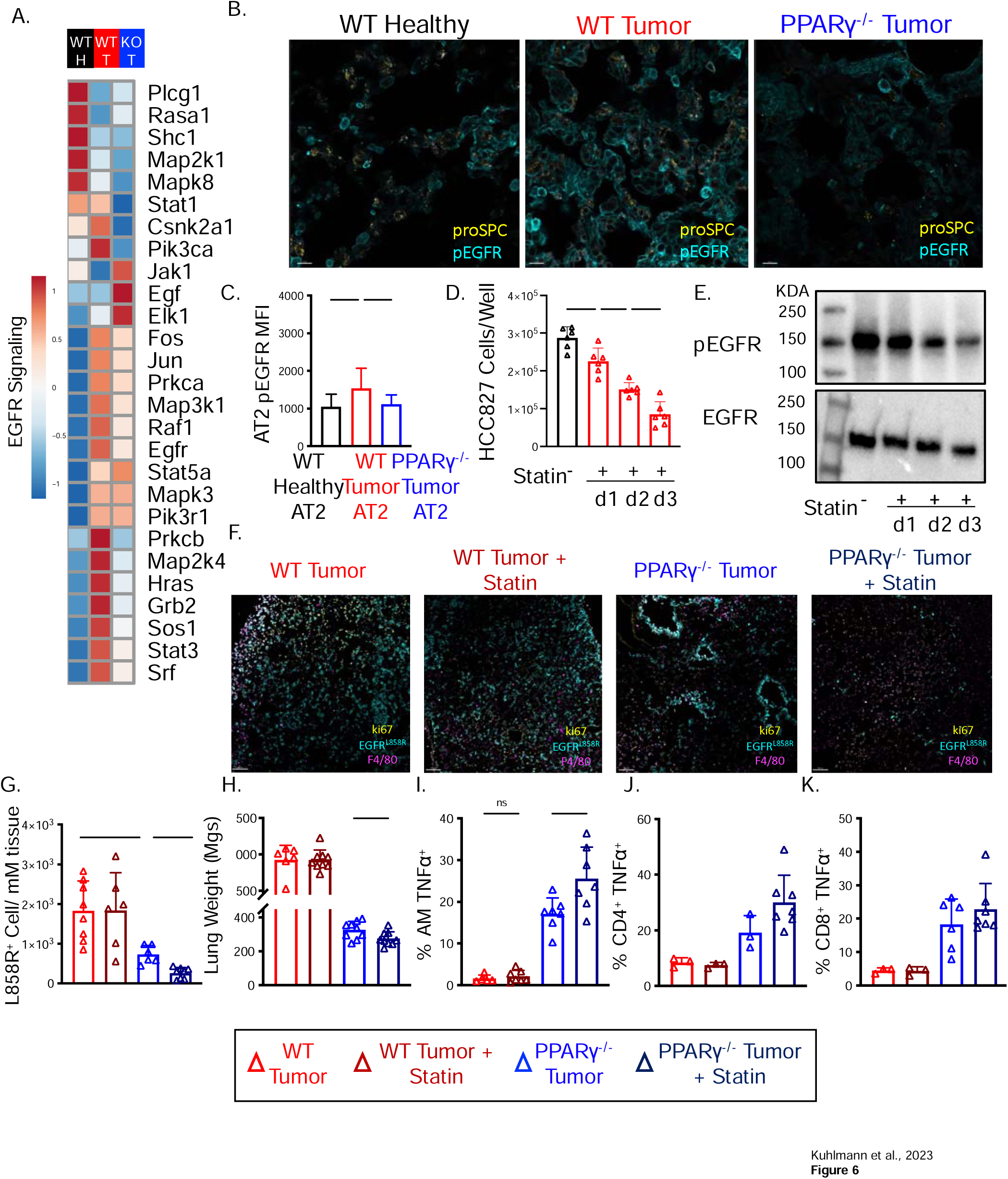
AM PPARγ rewires AT2 metabolism to support oncogenic signaling. (A) Heatmap shows expression of EGFR signaling genes by row Z-score (from scRNA-seq) in AT2 cells isolated from healthy lungs (black) or LUAD lungs that contain WT (red) or PPARγ TA-AMs (blue). (B-C) Lungs from same samples as in (A) were stained with mAbs to phospho-EGFR (pEGFR) and the AT2 cell marker pro-SPC and analyzed by confocal microscopy (B). Data are representative of sections taken from four separate mice. The intensity of pEGFR staining in AT2 was quantified using IMARIS (C). (D-E) The EGFR^Del19^ mutant human cell line HCC827 was cultured for 1-3 days in the absence or presence of 10 μM atorvastatin and each day the viable cell number was enumerated (D) and the amounts of pEGFR relative to total EGFR were assessed by Western blotting (E). (F-K) LUAD lungs containing WT (red) or PPARγ (blue) were treated by oral gavauge with pravastatin (0.5 mg, 5X/week) from weeks 3-7 on DOX. Tumor burden was measured using confocal microscopy staining with mAbs specific for the EGFR^L858R^ oncogene, macrophage marker F4/80, and ki67 (F, G) and by dry lung weight (H). (I-K) Intracellular cytokine staining, and flow cytometry was used to assess the frequency of TNF-producing AMs (I), CD4^+^ and CD8^+^ T cells (J, K) after stimulation with LPS (I) or PMA and ionomycin (J, K). Data shown are mean ± SEM, and statistical analysis were performed with a two-tailed unpaired Students test (E) or a two-way ANOVA (C,G,I-M). *p<0.05, **p < 0.01, ***p<0.001, ****p < 0.0001. Data are representative of ≥3 mice (B,C,F) and is pooled from ≥3 experiments with each group containing 50-149 cells (C) and 8 (D), 6-9 (G), 5-6 (H), 5-7 (I), 3-6 (J-K) mice.

Based off the clinical observations that LUAD patients fared better when treated with EGFR TKIs in combination with statin therapy to suppress cholesterol synthesis (53–55), we reasoned that the simultaneous decrease in EGFR-signaling and increase in cholesterol synthesis observed in AT2 cells in PPARγ^-/-^ Tumors would make them particularly susceptible to treatment with a statin. *In vitro* we replicated results (55) showing sensitivity of HCC827 cells to statins, specifically to atorvastatin (**Fig 6D**) but discovered this also causes a diminution in EGFR phosphorylation over time (**Fig 6E**). To test this idea *in vivo* we treated WT or PPARγ^-/-^Tumors with oral pravastatin from weeks 4-7 of tumorigenesis and then assessed tumor growth at week seven. While we did not see reduction in tumor burden in the WT Tumors treated with statin by either density of EGFR^L858R^-positive tumor cells by microscopy (**Fig 6F–G**) or lung weight (**Fig 6H**) we did see a significant reduction in the PPARγ^-/-^ Tumors treated with statins. Interestingly, the statin treatment also augmented the switch towards a more pro-inflammatory state of both AMs and T cells in the PPARγ^-/-^ Tumors relative to the WT Tumors (**Fig 6I–K**). These data indicate that the compensatory increase in cholesterol synthesis pathway by AT2 cells when TA-AMs are unable to metabolically support the tumor cells creates a metabolic vulnerability that can be therapeutically targeted with statins. In fact, this work identifies the entire GM-CSF—PPARγ—cholesterol efflux pathway as a novel and targetable axis, opening up several new alternative strategies for EGFR^+^ lung cancers.

## Discussion

The poor response rate to ICB and inevitability of resistance to targeted therapies against EGFR mutant LUAD leaves an unmet need for therapies that stimulate anti-tumor immunity beyond T cell centric modalities like PD-1/CTLA-4. TAMs are an ideal immunotherapeutic target because of their ubiquity and plasticity (6,7). TRMs, a subset of TAMs that have been increasingly implicated in tumorigenesis, are an intriguing candidate due to their unique capacity to meet the functional demand of their transformed client cells in a tissue-specific manner (33–36). Our study illuminates a novel mechanism by which malignant AT2 cells coopt homeostatic lipid metabolic functions of lung AMs to promote the formation of an immunologically permissive TME while rewiring macrophage metabolism to provide nutrients to sustain oncogenic signaling. Specifically, we found that tumor epithelium hijacks the GM-CSF-PPARγ signaling axis in AMs to obtain nutrients such as cholesterol. In the absence of this macrophage metabolic support, the tumor cells were forced to shift their own metabolic states to overcome and compensate for decreased nutrient supplementation, ultimately straining oncogenic EGFR signaling and presenting a targetable metabolic vulnerability in the tumor cells.

Our data identify a novel pro-tumor function of GM-CSF in the lungs by which it induces: (*i*) the accumulation of tumor-supportive AMs, (*ii*) PD-L1 expression and (*iii*) metabolic rewiring to support tumor metabolism. Constitutive expression of GM-CSF by AT2 cells-but not other sources-is required for AM development and maintenance (41). While the exact factors that lead to tonic GM-CSF secretion in AT2 cells is unknown, EGFR signaling induces AT2 cell GM-CSF secretion in allergy (56) and influenza (57), consistent with a well-established role for EGFR signaling inducing GM-CSF in keratinocytes (58). Thus, it is likely that constitutive oncogenic EGFR activity increases GM-CSF secretion in EGFR^+^ LUADs. In support of a positive feedback loop— in which epithelial GM-CSF ultimately promotes transformed AT2 cell proliferation via paracrine signaling with AMs— overexpression of GM-CSF in AT2 cells leads to their hyperplasia and accumulation of AMs specifically at loci with the most pronounced hyperplasia (42). Further, GM-CSF can also increase TAM production of hbEGF, an EGF-family member and growth factor (15,16). Collectively, these data support our model in which oncogenic EGFR signaling elevates GM-CSF that causes accumulation and PPARγ-dependent immunologic and metabolic reprogramming of TA-AMs, which ultimately fuels nutrients for oncogenic growth of tumor cells.

However, a contrasting role for GM-CSF in stimulating adaptive anti-cancer immune responses has also been described. While we reported an increase in PD-L1 expression on AMs following GM-CSF stimulation, the cytokine has an immunostimulatory effect on DCs (59). GVAX vaccines couple antigen presentation with local GM-CSF production and effectively promote anti-tumor immunity, but nonetheless have had lackluster performance in NSCLC patients (59). It remains to be explored under what scenarios and contexts GM-CSF can promote anti-tumor *vs.* protumor immune states.

Lipid accumulation in the TME is an emerging modality of immunosuppression (60–63). In our study, we explore how the tissue of tumor origin (i.e., lung) impacts the TME lipid composition and report increases in surfactant associated lipids (and proteins) in the BALF of tumor-burdened lungs in mice and humans. Both the lipid and protein component of surfactant have well-established immune-tolerizing roles (64) and coupled with the ability of GM-CSF to regulate phospholipid import and cholesterol efflux in AMs (50), we propose that the deregulated GM-CSF-surfactant homeostasis between AT2 cells and AMs is a central axis that has been previously underappreciated in how it controls immune function and metabolism in the TME and lung cancer progression.

Our data also suggest a model wherein ABCG1-mediated efflux of cholesterol from TA-AMs is transported to AT2 cells to support their EGFR signaling. Cholesterol is tightly controlled within the lung airways with more than 90% of the cholesterol in AT2 cells being imported (65) while secreted cholesterol only gradually exits into circulation via HDL (66,67). Given that ABCG1 is highly expressed by TA-AMs (in a PPARγ-dependent manner) and utilizes diverse cholesterol acceptors, including surfactant (66), we postulate that TA-AMs use ABCG1 to deliver cholesterol to tumor cells coupled to surfactant as part of the surfactant recycling process. It is also possible that there are other lipids, nutrients or metabolic supportive functions that the TA-AMs provide to the AT2 cells that our study may have missed. Additionally, ABCG1 expression has been shown to regulate M1 vs M2 macrophage skewing in ovarian cancer, bladder cancer and melanoma (67), suggesting it can also play an important role in the tolerogenic skewing of macrophages in multiple tumor types.

We have shown that rewiring AT2 cell cholesterol metabolism was detrimental for optimal oncogenic EGFR signaling. Cellular cholesterol content regulates EGFR with acute reduction in cholesterol content in lipid rafts promoting EGFR clustering and downstream signaling (68) but also promoting susceptibility to EGFR TKIs (69). Lung cancer patients receiving statins perform better on an EGFR targeted therapy (50). A Taiwanese study reported a significant reduction in death amongst patients receiving combo therapy with an impressive hazard ratio of 0.58 and an increased median survival of almost a full year (70). However, despite the high rate of EGFR mutations reported in this population (∼50%), the authors did not asses EGFR mutation status in this study and the specific interplay of EGFR and statins in the clinic remains to be determined. *In vitro*, EGFR mutant tumor cells that are treated with TKIs upregulate cholesterol synthesis (54). TKIs and statins synergize to induce apoptosis and delay the emergence of resistance, perhaps by impacting downstream farnesylation in proteins such as RAS by limiting cholesterol intermediates (53–55). We hypothesize that targeting AM metabolic support with a PPARγ inhibitor in addition to statin and TKI treatment is a promising future therapeutic target with the potential to inhibit oncogenic signaling and tumor growth while simultaneously engaging a potent anti-cancer immune response.

## Materials and Methods

### Mice

*Ccsp-rtTA; TetO-EGFR^L858R^* mice have been previously described (39). Pparγ^fl/fl^ ((71) mice crossed to Tg(Itgax^Cre^) were generously provided by Manfred Kopf (46). Pparγ^fl/fl^ ((71) mice were crossed to Tg(Csf1R^Cre^) JAX stock #029206 (72). SftpA/D^-/-^ mice were obtained from JAX, stock #027304 (73) (73). These mice were crossed to *Ccsp-rtTA;TetO-EGFR^L858R^* to create lung tumors harboring PPARγ deficient Alveolar Macrophages or lung tumors deficient in SFTAP/D. These mice were of an undefined mixed C57/BL6 and FVB background. Mice were fed a doxycycline containing chow (625 ppm) obtained from Harlan-Tekland. Both male and female mice were used at approximately equivalent levels and were randomly assigned to treatment groups. Only mice younger than three months old were induced. All animals were housed in a specific-pathogen-free facilities at the Salk Institute for biological sciences in accordance with the guidelines and regulations implanted by the Salk Institute Animal Care and use Committee or at Yale University in accordance with the Yale University’s Institutional Animal Care and Use Committee.

### Treatments

To deplete GM-CSF, mice were treated twice weekly via intraperitoneal (I.P.) injection of 0.5 mg anti-GM-CSF clone MP1-22E9 (BioXCell, Cat# BEO259) monoclonal antibody from BioXCell or IgG2a isotype control (Cat# BE0089). Treatments were initiated after two weeks of doxycycline administration and were continued for four weeks, at which point the mice were sacrificed and analyzed.

Mice were treated with 1mg/kg PPARγ antagonist GW9662 (Sigma Cat# M6191) to pharmacologically block signaling. Drug was suspended in corn oil and administered twice a week either intraperitoneally or intratracheally administration (I.T.). For I.T. administration, mice were briefly anesthetized with isoflurane before the trachea was exposed and no more than 30 uL of fluid was administered. These treatments commenced at two weeks of doxycycline and were continued for four weeks before the mice were sacrificed and analyzed.

Mice were treated with 0.5 mg of pravastatin (Caymen Cat# 10010342) dissolved in PBS which was given by oral gavage 5 times a week, starting at two to three weeks on doxycycline and continued for four weeks before the mice were sacrificed and analyzed.

### Bronchoalveolar lavage, alveolar macrophage and type II pneumocyte collection in mice

Epithelial lining fluid and non-adherent cells were collected from indicated mice by canulization of the trachea with a catheter. The lungs were then flushed three times with a 1LJml aliquot of PBS with a recovery of approximately 90% of fluid. The flushes were pooled and only samples with a consistent recovered volume were used in this study. The supernatant was removed and flash frozen in liquid nitrogen before storage at –80°C and the cellular pellets were resuspended in the culture media for isolation of alveolar macrophages by adherence to tissue culture plastic for at least 30 minutes.

Isolation of AT2 cells has been previously reported(74). Briefly mice were perfused using 10 mL PBS through the right ventricle of the heart. We then proceeded with tracheal canulization, and collection of BAL fluid as described above. The lungs were then inflated with 1 mL of elastase at 0.75 units/mL (Stemcell #07453) with 0.5 mg/mL DNAse (Sigma Cat# 10104159001) in HBSS (Gibco Cat# 13175-095). The trachea was then tied close using string and the inflated lungs were excised from the chest cavity. Lungs were washed with PBS (Life Tech Cat# 14190-144) and then placed in the incubator at 37°C for 10 minutes in a small round culture dish. After 10 minutes, lungs were finely chopped with a razor blade and an additional 1mL of digestion media was added to the plate which was returned to 37°C for an additional 5 minutes. After digestion, the contents of the plates were passed through a 70 uM filter which was also washed with 10 mL of RPMI-1640 (Life Tech Cat# 11875-093) media plus 10% FBS (Omega Scientific Cat# FB-02). After centrifugation, cell pellets were lysed with ACK lysis buffer (KD Medical RGF-3015), washed, and then re-pelleted. Beads were then used to further purify AT2 cells. Briefly, cell pellets were resuspended in MACs buffer (2% FBS in PBS + EDTA (Life Tech Cat# 15575-038)) and Biotin-selection cocktail (Stem Cell Cat# 17683) was used to negatively select against CD45 and CD31. Following negative selection, cells in the flow through were then positively selected against EPCAM using PE-selection cocktail (Stem Cell Cat# 17684) and a population of approximately 90% purity was then obtained.

### Magnetic Resonance Imaging

Magnetic resonance images were collected using a mini-4T horizontal-bore spectrometer (Bruker AVANCE). During data collection animals were anesthetized with a constant flow of isofluorane and oxygen (2-2.5% v/v). Tumor burden was subsequently quantified for each animal by calculating the volume of visible lung opacity within the chest cavity in every image in the stack using the BioImage Suite software (75).

### Cell lines

The female human lung adenocarcinoma cell line HCC827 with an acquired exon-19 deletion (ΔE746-A750) in EGFR’s tyrosine kinase domain (ATTC Cat# CRL-2868, RRID:CVCL_2063) were obtained from the Leibel lab and were used for less than 15 passages. Cells were cultured in RPMI-1640 media supplemented with 10% fetal bovine serum (FBS) and 1% penicillin-streptomycin (pen/strep) (Life Tech #15140-122). To generate 3-D lung spheroids, HCC827 cells were dissociated using TrypLE Express media (Thermo Scientific Cat# 12604021) counted, spun down and subsequently resuspended in RPMI+10% FBS + 2% Pen/Strep media. 10,000 cells per well were plated in ultra-low attachment 96 well plates (Sigma Cat# CLS3474). Media was replenished on d3 and d7, at which point spheroids were formed. For co-culture experiments, 50,000 ex-vivo alveolar macrophages were added to wells on d7 with 10 ng of recombinant mouse GM-CSF (R&D Cat# BJ2520011) added daily. Spheroids were dissociated using Acutase (Stem Cell Cat# 07920) after 3 days of co-culture and single cell suspensions were analyzed by flow cytometry using surface and intracellular staining as described below.

### Human BALF Samples

This protocol was approved by the IRB at UCSD biorepository. Each subject gave written, informed consent prior to enrollment in this study. Both male and female patients were included and features such as age and sex are depicted in table S1. Inclusion criteria were based off of diagnosis with lung cancer, mycobacterial infection or COPD. Exclusion criteria included patients who had more than one diagnosis at a time, though patients with neoplasms at alternate sites were included. Samples were collected after instillation of sterile saline with attempt for 50% or greater return from the targeted lung lobe. No complications occurred with sample collection during the procedure. For processing, samples were first filtered over 150 uM nylon mesh to remove debris and mucus. Cells were pelleted at 500g and the fluid was recovered and frozen in liquid nitrogen before storage at –80°C. RBCs were lysed from the cell pellet using ACK lysis buffer. Cells were then counted and frozen in FBS containing 10% DMSO (Fisher/Corning MT25950CQC) and stored in liquid nitrogen until further analysis.

### Lipid measurements using LC/MS

Lipid extraction from cells or BALF was performed using a modified Bligh-Dyer method (76) and as previously reported (63). Cell pellets were diluted in 1 mL PBS and they or BALF were shaken in a glass vial (VWR) with 2 mL chloroform and 1 mL methanol containing internal standards (^13^C_16_-palmitic acid, d7-Cholesterol) for 30 seconds. Vials were then vortexed for 15 seconds and centrifuged at 2400 x g for 6 mins during which phases separation occurred. A Pasteur pipette was used to retrieve the bottom (organic) layer, which was then dried under a gentle stream of nitrogen, and reconstituted in 2:1 chloroform:methanol for LC/MS analysis. A Vanquish HPLC online with a Q-Exactive quadrupole-orbitrap mass spectrometer equipped with an electrospray ion source (Thermo) was used to lipidomic analysis. Solvent A contained of 95:5 water:methanol, solvent B consisted of 60:35:5 <u>isopropanol</u>:methanol:water. For acquisition of data in positive ionization mode, solvents A and B consisted of 5 mM ammonium formate with 0.1% formic acid; for data acquisition in the negative ionization mode, solvents contained 0.028% ammonium hydroxide. A Bio-Bond (Dikma) C4 column (5 μm, 4.6 mm × 50 mm) was used. The gradient was held at 0% B between 0 and 5 min, subsequently raised to 20% B at 5.1 min, then increased linearly from 20% to 100% B between 5.1 and 55 min, before being held at 100% B between 55 min and 63 min, before being returned to 0% B at 63.1 min, and then held at 0% B until 70 min. Flow rate was 0.1 mL/min from 0 to 5 min, 0.4 mL/min between 5.1 min and 55 min, and 0.5 mL/min between 55 min and 70 min. Spray voltage was 3.5 kV for positive ion mode and 2.5 kV for negative ionization mode. Sheath, sweep, and auxiliary, and gases were 53, 3 and 14, respectively. Capillary temperature was kept at 275°C. Data was collected in full MS/dd-MS2 (top 5) mode. Full MS was attained from 100–1500 m/z with resolution of 70,000, AGC target of 1 × 10^6^ and a maximum injection time of 100 ms. MS2 was acquired with resolution of 17,500, a fixed first mass of 50 m/z, AGC target of 1 × 10^5^ and a maximum injection time of 200 ms. Stepped normalized collision energies were 20, 30 and 40%. LipidSearch was performed to identify lipids. Mass accuracy peak integration, and chromotography, of all LipidSearch-identified lipids were verified using Skyline (77). Peak areas were used in data reporting, and data was normalized using internal standards.

### Protein quantification and enzyme-linked immunosorbent assays (ELISAS) in Mouse and Human BALF

Quantification of total protein levels from BAL fluid was measured using a BCA protein assay kit (Thermo Fisher Cat# 23227) according to the manufacturer’s instructions. A sandwich ELISA was used to quantify mouse SP-D (R&D Cat# DY6839-05) and human SP-D (R&D Cat# DY1920) using R&D duoSet antibody pairs. Wells were coated using the manufacturers recommended concentration of capture antibody followed by detection with a biotinylated monoclonal antibody and then the colorimetric changes were quantified. Pre-coated sandwich ELISA plates were used to quantify mouse SP-A (Novus Biologicals Cat# NBP2-76693).

### Tumor digestion and cell isolation

Following BALF isolation, lungs were excised from the chest, washed with PBS, dried on paper towels and then weighed. Lungs were minced into small pieces and suspended in digestion media consisting of RPMI 1640 with 2% FBS, 0.5 ug/mL of DNase I and 1 unit/mL collagenase Type I (Sigma-Aldrich Cat# 0000137295) and placed in an incubator at 37°C for 45 minutes, shaking once during digestion. Lungs were then mashed against 70 μM cell strainers (VWR Cat# 10199-657) to filter and then red blood cells were lysed using ACK lysis buffer, mixed with RPMI 1640 containing 10% FBS and 1% pen-strep, centrifuged at 400g for 5 minutes at 4°C to obtain a single-cell suspension.

### Uptake of fatty acids, cholesterol, or phospholipids and neutral lipid content assay

Cells were incubated in PBS containing 0.5 μg/ml C1-BODIPY® 500/510 C12 (ThermoFisher Cat# D3823), PBS containing 10uM NBD Cholesterol (Cayman #30136), or 0.25 μg/ml DPPE-PE (Avanti Cat# 810158P) for 30 minutes at 37°C in order to measure the uptake of fatty acids or cholesterol bodipy phospholipids. Cells were washed after incubation with MACS buffer (PBS containing 2% FBS) to achieve surface staining.

### Flow Cytometry, cell sorting, and antibodies

Prior to staining, single cell suspensions were incubated on ice for 10 minutes with Fc receptor-blocking anti-CD16/32 (BioLegend Cat# 101301). Cell suspensions were first stained for 5 minutes at room temperature with LIVE/DEAD® Fixable Red Dead Cell Stain Kit (ThermoFisher Cat# L23102). Surface proteins were then stained for 30 minutes at 4°C in FACS buffer (PBS containing 2% FBS and 0.1% NaN3 (MP Bio Cat# 2102891-CF)). Cell suspensions were re-suspended in RPMI 1640 containing 10% FBS, itself stimulated by 50 ng/ml PMA (Phorbol 12-myristate 13-acetate) (Sigma Cat# P8139) and 3 μM Ionomycin (Sigma Cat# 10634) in the presence 2.5 μg/ml Brefeldin A (BioLegend Cat# 420601) for 4 hours at 37°C to detect cytokine production *ex-vivo*. For surface marker staining, cells were processed as described above. Cells were fixed in BD Cytofix/Cytoperm (BD Cat# 554714) for 30 minutes at 4°C, then washed with 1 × Permeabilization buffer (Invitrogen #00-8333-56) to measure intracellular cytokine staining. Cells were fixed in Foxp3 / Transcription Factor Fixation/Permeabilization buffer (Invitrogen Cat# 00-5521-00) for 30 minutes at 4°C, then washed with 1 x Permeabilization buffer (three times) to assess nuclear protein staining. Subsequently, cells were stained for 30 minutes at 4°C with intracellular antibodies. The LSR-II flow cytometer (BD Biosciences) was employed to process samples, and FlowJo V10 (TreeStar RRID:SCR_008520) was used to analyze data. Either the FACSAria III sorter or Fusion sorter (BD Biosciences) were used to sort cells.

The following list of antibodies and their concentration against mouse proteins were employed: αCD45 (1:400; BioLegend Cat# 103147, RRID:AB 2564383), αCD3ε (1:300; Thermo Fisher Scientific Cat# 46-0032-82, RRID:AB_1834427), αCD4 (1:300; BioLegend Cat# 100406, RRID:AB_312691), αCD8a (1:300; BD Biosciences Cat# 741811, RRID:AB_2871149), αCD11b (1:800; BD Biosciences Cat# 612801, RRID:AB_2870128), αLy6C (1:300; BD Biosciences Cat# 563011, RRID:AB_2737949), αLy6G (1:300; BioLegend Cat# 127624, RRID:AB_10640819), αCD11c(1:300; BD Biosciences Cat# 564080, RRID:AB_2738580), αSiglecF (1:300; BD Biosciences Cat# 746668, RRID:AB_2743940), αTNFα (1:300; (BioLegend Cat# 506305, RRID:AB_315426), αCD200R (1:300; BioLegend Cat# 123908, RRID:AB_2074080), αPD-L1 (1:300; BioLegend Cat# 124321, RRID:AB_2563635), αKi67 (1:300; (Thermo Fisher Scientific Cat# 56-5698-82, RRID:AB_2637480), αGM-CSFR (1:300; Thermo Fisher Scientific Cat# MA5-23918, RRID:AB_2608189), αGM-CSF(1:100; BioLegend Cat# 505406, RRID:AB_315382), αMHCII (1:300; BioLegend Cat# 107624, RRID:AB_2191073), αABCG1 (1:200; Bioss Cat# bs-1231R-FITC, RRID:AB_11120127), αEPCAM (1:300; BioLegend Cat# 118212, RRID:AB_1134101), αCD31 (1:300; MEC13.3, BioLegend Cat# 102424, RRID:AB_2650892) and αproSPC (1:300; Abcam Cat# ab270521).The following antibody and the concentration used against human proteins were employed: αEGFR (1:1000 for western blot; Cell Signaling Technology Cat# 4267, RRID:AB_2246311), αEGFR^L858R^ mutant specific (1:100 microscopy; Cell Signaling Technology Cat# 3197, RRID:AB_1903955), αpEGFR (1:100; phospho Y1069) (Abcam Cat# ab205828, RRID:AB_2890267).

### Isotopic Tracing of TCA cycle intermediates, FAO

Metabolites were extracted using a modified Bligh and Dyer method and analyzed as previously described in detail [65]. Cells were cultured in DMEM (Sigma-Aldrich, St. Louis, MO, USA, Cat# 5030) medium containing 25 mM [U-^13^C_6_]glucose. (Cambridge Isotopes Inc. (Tewksbury, MA, USA), Cat# CLM-1396-25). To limit processing time, AMs were directly isolated from BALF, counted, and then plated in standard DMEM+10% FCS media for 2 hours during which cells adhered to the bottom of the plate, after which cells were washed and tracer medium was added. For glucose isotopic labeling experiments, cells were cultured in tracer medium for four hours. Labeling (corrected for natural abundance using in-house software) is depicted as labeled fraction of metabolites (mole percent enrichment (MPE) times abundance).

For isotopic tracing assessment of FAO and import, cells were isolated from BALF as described above and cultured with [U-^13^C_16_]palmitate (Cambridge Isotopes Inc. (Tewksbury, MA, USA), Cat# CLM-409-PK) for 6 hours in DMEM containing 10% delipidated FBS before reaction was stopped with ice cold methanol. [U-^13^C_16_]palmitate was noncovalently bound to fatty acid-free BSA and added to culture medium at 5% of the final volume (50 μM final concentration). Lipids were extracted as described by others(78) and M2 labelling on citrate was used to determine rates of fatty acid oxidation. Analysis of the percentage of [U-^13^C_16_]palmitate that comprised the total intracellular pool of palmitate was used to look at lipid import. Calculation of isotopologue distribution has been previously described in great detail (78).

### Radioactive isotopic Fatty Acid Oxidation and Sterol Synthesis and Fatty Acid Synthesis Assays

Use of radioactive palmitate to measure FAO in immune cells has been previously described (63). Briefly, cells were isolated ex-vivo and then incubated for six hours with 0.5 μCi ^14^C-palmitic acid (Perkin Elmer Cat# NEC075H050UC) in the following media: RPMI 1640 medium containing 2% FBS, 10 μM palmitic acid, 1% fatty acid free BSA (Sigma Cat# A3294), 500 μM carnitine (Sigma Cat# C0283). Oxidation was stopped by addition of ice-cold methanol, lipids were precipitated with 10% trichloroacetic acid, and subsequently loaded into ion exchange columns containing DOWEX 1X2-400 resin (Sigma). FAO product was eluted with water and radiation was measured by liquid scintillation.

For fatty acid and sterol synthesis, cells were labeled with 0.5 uCi of 56 mCi/mmol [^14^C] acetic acid (Perkin Elmer Cat# NEC084H001MC) per well overnight. Cells were then washed with PBS twice, lysed in 100 ul 0.2 M KOH, supplemented with 60ul 50% KOH and 400 ul EtOH. The samples were then incubated at 75°C for 2 hours, then at room temperature for 2 hours. Sterols were extracted by adding 900 ul hexane (organic phase). The remaining aqueous phase was acidified by adding 100 ul 12M HCl. Fatty acids were extracted by adding 900 ul hexane (organic phase). Both samples for sterols and fatty acids were dried and resuspended in 1:1 chloroform and hexane before sending to scintillation counting.

### Cholesterol Esterification and Cholesterol Efflux assays

Cholesterol efflux was measured using a cell-based cholesterol efflux kit (BioVision Cat# K582-100) in accordance with the manufacturer’s instructions. AMs purified from mouse BALF were plated at a density of 2×10˄4 and allowed to adhere to the plate. Cells were then incubated with fluorescent cholesterol for an hour, washed, and allowed to equilibrate overnight. FBS in phenol red free RPMI was then added, and efflux was carried out over the ensuing four hours after which timepoint the supernatant was removed, and cellular contents were lysed. Efflux rate was calculated as the amount of cholesterol in the supernatant divided by the sum of the signal from the supernatant and intracellular compartment. 10 ng/mL GM-CSF and 10ug/mL SOAT1i Sandoz 58-035 (Sigma Cat# S9318-5MG) were added to the media during equilibration to assess their impact on efflux rates

### Seahorse-Basal Mitochondrial Respiration

Alveolar Macrophages purified from BALF was plated in XF media (Agilent Cat# 103681-100) containing glucose, L-glutamine, and sodium pyruvate were plated at a density of 2×10˄4 cells per well. Oxygen consumption rate (OCAR) and extracellular acidification rate (ECAR) were measured using an Agilent Seahorse DF396 Analyzer system (Seahorse Biosciences) in accordance with the manufactures instructions.

### Previously published bulk RNA-Seq analysis

We analyzed expression data from previously published and publicly available bulk RNA-sequencing data from our lab (39). Details of sample collection, sequencing, and data processing are described in detail elsewhere (39).

### Single-cell RNA sequencing

We sorted four distinct populations of cells (AMs: CD45^+^CD11c^+^SigF^+^, IMs: CD45^+^CD11b^+^Ly6G-, T cells: CD45^+^CD3^+^ and epithelial cells: CD45-EPCAM^+^ from healthy, tumor-bearing, and PPARγ^-/-^ tumor-bearing lungs. An equal number of each group of cells were partitioned into 10x Genomics Chromium Single Cell Controller chips to form an emulsion of nano-liter-sized droplets. Chromium Single Cell 3’ library and gel bead kit v2 (10X Genomics, Cat# PN-120237) was used to create RNA sequencing libraries. We loaded droplets comprising individual cells, RT reagents encapsulated in a gel bead and loaded it with poly(dT) primers which included a 16 base cell barcode and an additional 10 base UMI. Following reverse transcription reactions

A barcode of full-length cDNA was created following RT reactions which was cleaned up suing recovery agent and DynaBeads MyOne silane Beads (Thermo Fisher Scientific, Cat# 37002D). cDNA was then amplified, Chromium Single Cell 3′ v2 Reagent Kit was used to construct an indexed sequencing library. A NextSeq500 Sequencing System (Illumina Cambridge) was used to sequence the libraries.

### Single-cell RNA sequencing data processing and analysis

Pipeline analysis that we used has been previously described (63). Briefly, we processed output files using Cell Ranger (version 2.1.1) to generate a fastq file that had been aligned to the mouse genome. The Seurat package (version 3.1.5) in RStudio was used for further analysis. Prior to analysis, we used the following filter parameters: a gene must be expressed in greater than 3 cells, and to avoid low quality or doublets, cells uniquely expressed genes must fall within the range of 200-3,000 genes, and any cell with greater than 8% of tis reads mapping to mitochondrial genes was discarded to account for dying cells. Following quality control, we normalized our gene expression matrix and subsequently natural log-transformed it to identify highly variable features. We then integrated samples from Healthy, WT, and AM PPARγ^-/-^tumors to obtain a single expression matrix comprising genes from 6,060 cells (1329 Health cells, 2,292 WT Tumor cells, 2479 PPARγ^-/-^tumor cells). Heatmaps were calculated using the pheatmap function, and DEGs were found using the FindMarkers in the Seurat package. Single-cell gene set enrichment analysis was done using R package “singleseqgset” (https://github.com/arc85/singleseqgset). For gene expression heatmaps, average expression matrices of all clusters were extracted from the Seurat object and plotted using R package “pheatmap” representing z-scores of each row. The analysis of different tumor macrophages from a publicly available gene set (GSE97168) (47) was performed similarly to the above-described workflow. “Enrichment score” of hallmark gene sets were calculated using Seurat’s “AddModuleScore()” function.

### Immunofluorescence microscopy

Euthanized mice were perfused with 30 mL PBS using a cardiac puncture. Lungs were dissected at the stated time post tumor initiation and fixed for 24 hours in 2% paraformaldehyde (Santa Cruz Biotechnology Cat# sc-253236) at 4°C. The lungs were then equilibrated in 30% sucrose for two days at 4°C and embedded in OCT medium which was frozen in 2-methylbutane using liquid nitrogen and then sectioned into 12μm sections using a cryostat. The tissues were kept at –80°C. Prior to staining with antibodies of interest using a free-floating method, the sections were thawed at room temperature for 10 mins then washed with PBS for 5 mins at room temperature. Prior to staining with antibodies of interest, the sections were blocked in Blocking Buffer Buffer (1% BSA) and 0.05% NaN3 in PBS with the addition of Fc blocking antibody to prevent non-specific binding for 2 hours at room temperature then washed three times in Wash Buffer (0.1% of Tween-20 (Thermo Fisher Scientific Cat# 28321) in PBS). Primary antibodies were diluted in Staining Buffer (2% BSA, 0.01% NaN3, and 0.5% Tween 20) and added to free floating sections for 24-48 hours at 4°C. Tissues were washed twice in Washing Buffer then secondary antibodies were diluted in Staining Buffer and added to free floating sections for 24 hours at 4°C. The tissues were washed twice in Washing Buffer, mounted onto Epredia Polysine Slides (Fisher 12-545-78) using prolong gold antifade mounting reagent (Thermo Fisher Cat# P36934) then covered with No.25 Glass Coverslips (Epredia Cat# 24×50-1.5-001G).

Images were collected on a Zeiss LSM 880 Rear Port Laser Scanning Confocal Microscope with or without Airyscan FAST module using either 10× or 20× air objectives. To visualize tumor cell density, images of mice induced for 8-9 weeks and or 8-9 weeks of induction with 3 weeks of statin treatment were collected using the 10× objective with 0.7 zoom. To quantify the density of cells, the images were analyzed in Imaris (software version 9.9.0, RRID:SCR_007370). In Imaris, EGFR^L858R^ cells were individually defined as surfaces and counted. Density was determined following normalization to the area quantified. This process was conducted in 2 total regions from 3-4 mice.To visualize pEGFR intensity, images of mice induced for 8-9 weeks were collected using the 20× objective with 0.7 zoom. To quantify the density of cells, the images were analyzed in Imaris software version 9.9.0. In Imaris, proSPC^+^ cells were individually defined as surfaces. Then, the “mean fluorescence intensity” function for the chanel containing pEGFR in Imaris was applied to quantify intensity. This process was conducted in a total of 2 regions of interest from 3 different mice.

### Statistical Analysis

Statistical analyses were performed in GraphPad Prism (v9.2.0, RRID:SCR_002798). One-way ANOVA with multiple comparison testing, or two-tailed paired or unpaired Students *t*-tests were used to determine statistical significance (\**P* < 0.05, \*\**P* < 0.01, \*\*\**P* < 0.001, \*\*\*\**P* < 0.0001) as indicated in the figures.

## Acknowledgements

We thank all members of the Kaech and Politi labs for helpful discussions. We would like to thank Carolyn O’Connor of the Salk Flow Cytometry Core, April Williams of the Salk Bioinformatics core and Antonio MF Pinto of the Salk Mass Spectrometry Core. Pparγ^fl/fl^Cd11c^Cre^ were generously gifted by Dr. Manfred Kopf. This work was supported by National Institute of Health (NIH) RO1 CA230275, and RO1CA195720 to S.M.K. and K.P., by the Mark Foundation for Cancer Research and by P30CA014195 to S.M.K. and R.S. AKH was supported by the Yale University Interdisciplinary Immunology Training Grant (T32 AI-007019), Yale Cancer Biology Training Program (T32 CA193200-01A1) and by a diversity supplement for National Institute of Health (NIH) RO1 CA216101S1. This work was supported by the Flow Cytometry Core Facility at Yale University and at the Salk Institute with funding from the NIH-NCI CCSG:P30 014195 and Shared Instruments Grant S10-OD023689 (Aria Fusion cell sorter). This work was further supported by the Waitt Advanced Biophotonics Core Facility of the Salk Institute with funding from NIH-NCI CCSG: P30 014195 and the Waitt Foundation. The NGS Core Facility of the Salk Institute is supported by NIH-NCI CCSG: P30 014195, the Chapman Foundation and the Helmsley Charitable Trust. The Mass Spectrometry Core of the Salk Institute is supported by funding from NIH-NCI CCSG P30 014195 and the Helmsley Center for Genomic Medicine. The MS data were gathered on a ThermoFisher Q Exactive Hybrid Quadrupole Orbitrap mass spectrometer funded by NIH grant 1S10OD021815001.

## Author Contributions

AKH, SMK and KP conceptualized, designed, and supervised the research. AKH performed experiments with assistance from TC, ZX, KAT, CRO, SRL and EMK. AKH analyzed the generated data. TC performed flux tracing experiments and ZX performed radio-isotopic tracing experiments. MN and GZC provided clinical samples. RS, SLL, and CMM provided scientific input. AKH and SMK prepared the manuscript.

