## Supplemental Figures 1-4 and Supplemental Table 1 for "EGFR^+^ lung adenocarcinomas coopt alveolar macrophage metabolism and function to support EGFR signaling and growth"

### Fig S1: Myeloid cell regulation during lung tumorigenesis.

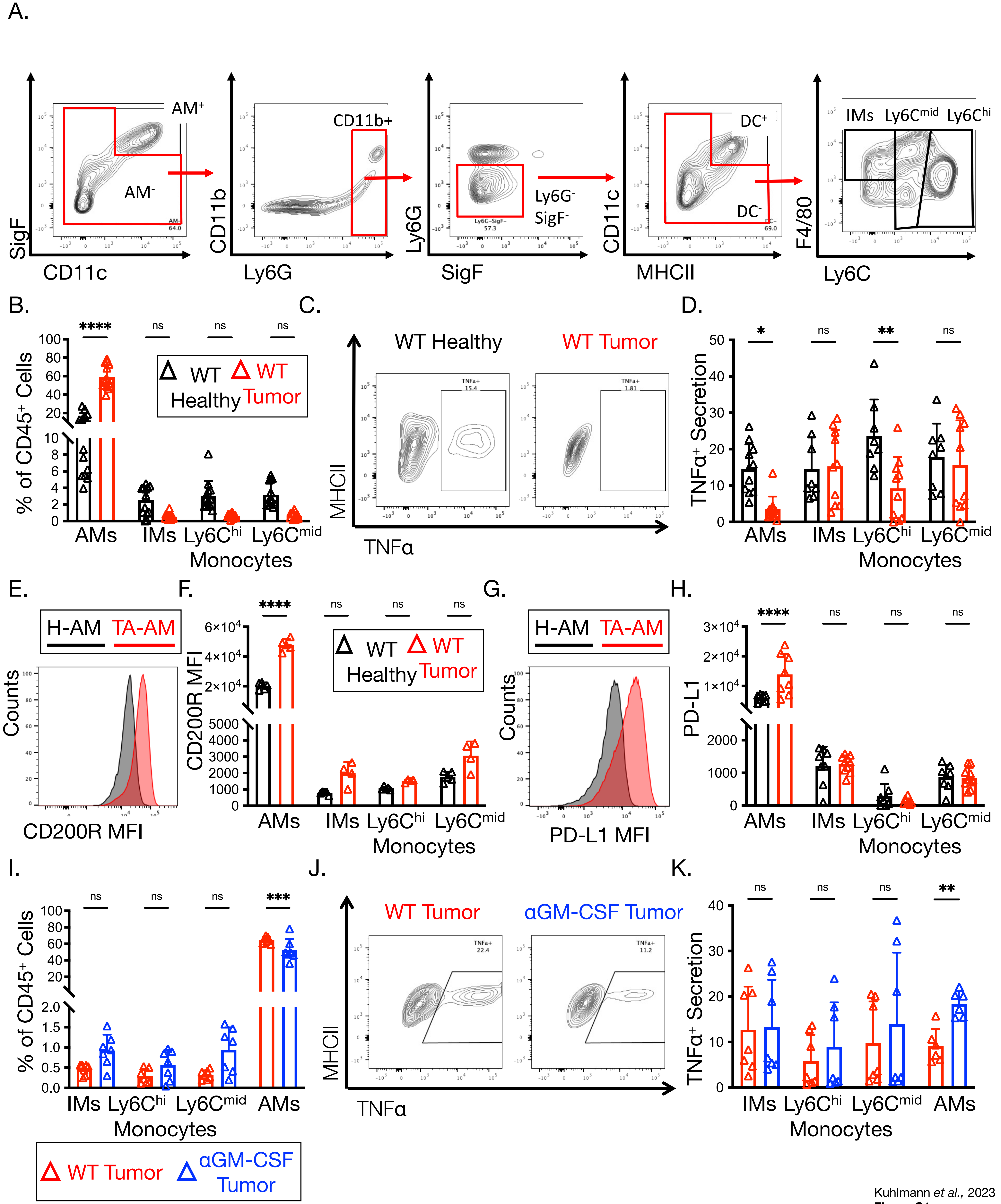


Kuhlmann *et al.*, 2023
**Figure S1**

**Figure S1. Myeloid cell regulation during lung tumorigenesis.**

(**A**) Flow cytometry plots of representative gating on myeloid cells within the TME: AMs- (CD45$^{+}$SigF$^{+}$CD11c$^{+}$), CD11b$^{+}$DCs$^{-}$ (CD45$^{+}$AM$^{-}$CD11b$^{+}$Ly6G$^{-}$SigF$^{-}$CD11c$^{+}$MHCII$^{hi}$), IMs- (CD45$^{+}$AM$^{-}$CD11b$^{+}$Ly6G$^{-}$SigF$^{-}$DC$^{-}$F4/80$^{+}$), Ly6C$^{mid}$ monocytes- (CD45$^{+}$AM$^{-}$CD11b$^{+}$Ly6G$^{-}$SigF$^{-}$DC$^{-}$Ly6C$^{mid}$), Ly6C$^{hi}$ monocytes- (CD45$^{+}$AM$^{-}$CD11b$^{+}$Ly6G$^{-}$SigF$^{-}$DC$^{-}$Ly6C$^{hi}$). **(B-H)** After 6-8 weeks on DOX, myeloid cell infiltrates from littermate control (WT Heathy, black) mice or those with fully established LUAD (WT Tumor, red) were quantified **(B),** stimulated *ex vivo* with LPS to measure TNF production **(C-D)**, or left unstimulated and assessed for expression of the inhibitory receptor CD200R **(E-F)** and checkpoint ligand PD-L1 **(G-H)**. **(I-K)** GM-CSF blocking antibodies were administered to LUAD mice i.p. (0.5mg/mouse) 2x a week between weeks 4-7 on DOX. Following GM-CSF depletion, **(I)** the number of myeloid cells present in the lungs was quantified and **(J-K)** cells were stimulated with LPS to measure TNF secretion.

Data shown are mean ± SEM, and statistical analysis were performed with a two way ANOVA. *p<0.05, **p < 0.01, ***p<0.001, ****p < 0.0001. Data depicts representative histograms (E,G) or flow plots (J) from flow cytometer. Data are pooled from ≥3 experiments with each group containing 13-14 (B), 8-10 (D,F), 5-4 (H), or 7(I,K) mice.

### Fig S2: Lipid import from myeloid cells in the TME

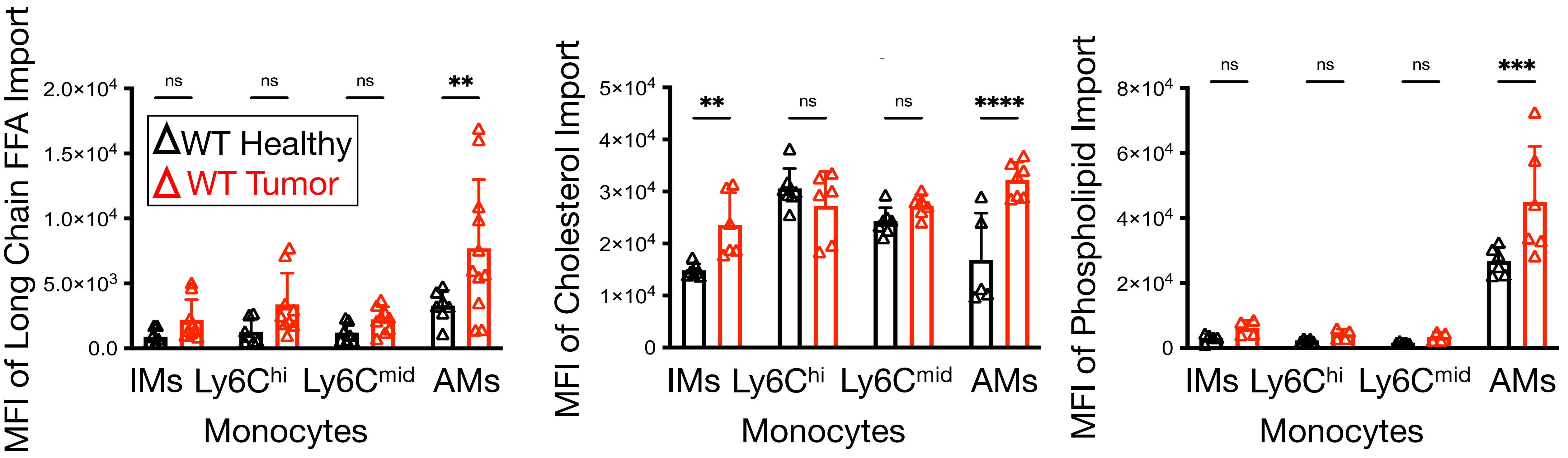


Kuhlmann et al., 2023
**Figure S2**

**Figure S2. Lipid import from myeloid cells in the TME**

Import of free fatty acids, cholesterol and phospholipids (Bodipy C12, NBD cholesterol, DPPE respectively) were compared between myeloid cells isolated from lungs of littermate controls (WT Healthy, black) and LUAD mice (WT-Tumor, red) using flow cytometry. MFI is shown in the bar graphs.

Data shown are mean ± SEM, and statistical analysis were performed by two way ANOVA. *p<0.05, **p < 0.01, ***p<0.001, ****p < 0.0001. Data are r pooled from ≥2 experiments with each group containing 4-9 animals.

### Fig S3 Macrophage or Dendritic Cell Deficiency of PPARγ alters the tumor immune microenvironment.

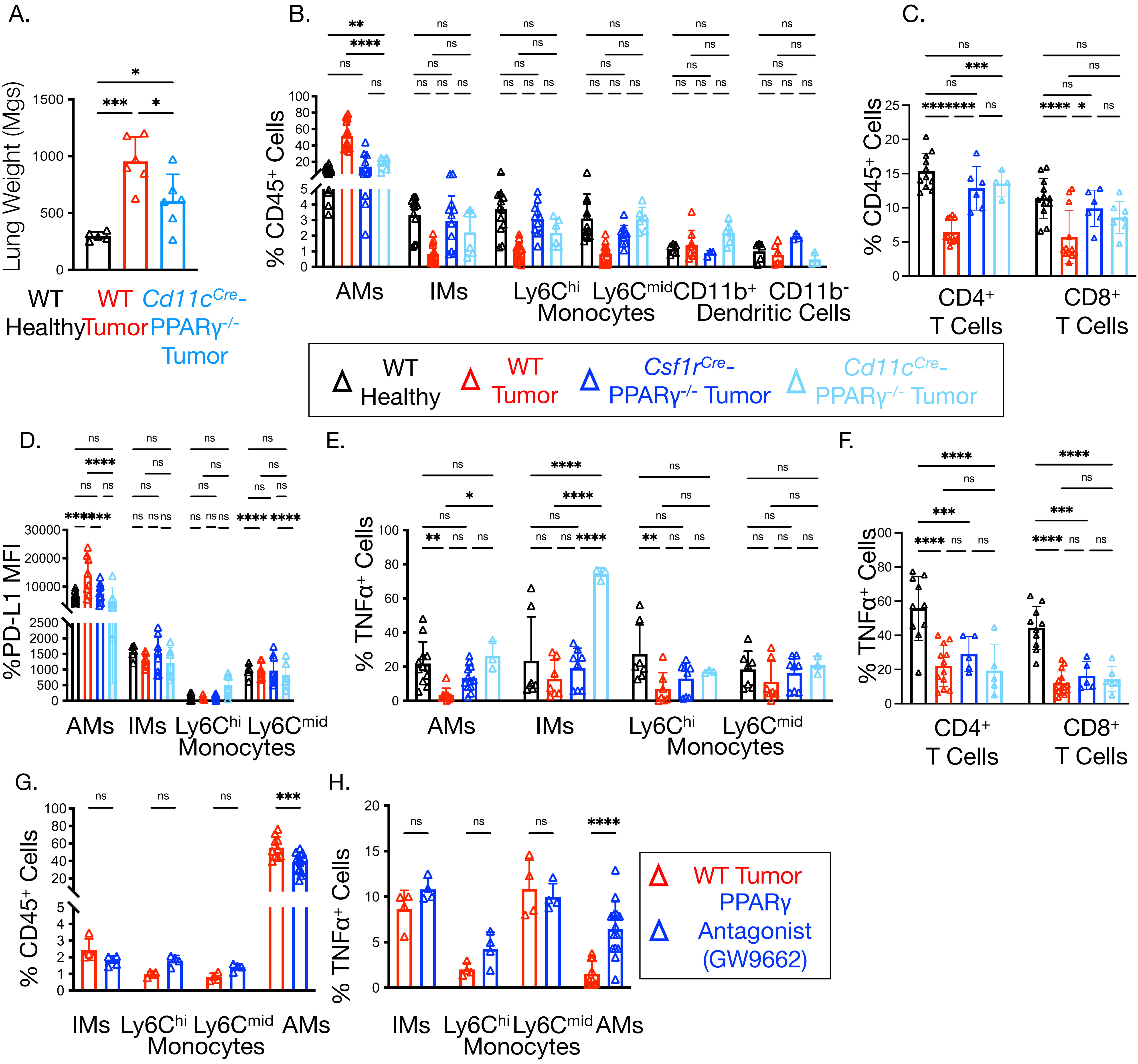


Kuhlmann *et al.*, 2023
**Figure S3**

**Fig S3 Deficiency of PPARγ in CSF1R- or CD11c-expressing cells suppresses tumor growth.** LUAD mice with deletion of PPARγ in macrophages (*Pparγ$^{Fl/Fl}$; Csf1r$^{Cre}$, Ccsp-rtTA; TetO-EGFR$^{L858R}$* (dark blue)) or in dendritic cells and AMs (*Pparγ$^{Fl/Fl}$Cd11c$^{Cre}$, Ccsp-rtTA; TetO-EGFR$^{L858R}$* (light blue)) and their LUAD littermate 'WT Tumor' controls (*Pparγ$^{Fl/F}$; Ccsp-rtTA; TetO-EGFR$^{L858R}$* (red)) along with 'WT Healthy' controls (*TetO-EGFR$^{L858R}$* or *Ccsp-rtTA* mice (black)) were placed on DOX for 7-9 weeks and then tumor immune infiltrates were examined by flow cytometry. **(A)** Bar graphs show the dry lung weights. **(B-C)** Bar graphs show frequency of myeloid cells (AMs, immature AMs, IMs, monocytes, and cDCs) **(B)** or of T cells **(C)** The following gating strategy was used: AMs: $CD45^+CD11b^-SigF^+CD11c^+$; immature AMs: $CD45^+CD11b^+SigF^+CD11c^+$; IMs: $CD45^+AM^-CD11b^+Ly6G^-$; DCs: $CD45^+CD11c^+MHCII^{hi}SigF^-Ly6G^-$; $CD4^+$ and $CD8^+$ T cells were gated on $CD45^+CD3^+$ cells. **(D)** MFI of PD-L1 expression on AMs, IMs, and monocytes. **(E-F)** Percentage of TNF-secreting myeloid cells following LPS stimulation **(E)** or T cells following PMA/ionomycin stimulation **(F)**. **(G-H)** LUAD mice and littermate controls were placed on DOX for four weeks and then treated by oral gauvage 5x/week with the PPARγ antagonist GW9662 (1mg/kg) in corn oil or vehicle alone for three more weeks. Bar graphs show frequency of myeloid subsets (AMs, IMs, monocytes, and cDCs) **(G)** and percentage of those that produce TNF after LPS stimulation **(H)**.

Data shown are mean ± SEM, and statistical analysis were performed with a two-way ANOVA. *p<0.05, **p < 0.01, ***p<0.001, ****p < 0.0001. Data are pooled from ≥2 experiments with each group containing 6 (A), 5-15(B), (4-12 (C), 6-10 (D), 3-12 (E), 5-14 (F), 3-6 (G) and 4-12 (H) mice.

### Fig S4: Tumorigenesis rewires LUAD patient macrophage lipid and mitochondrial metabolism.

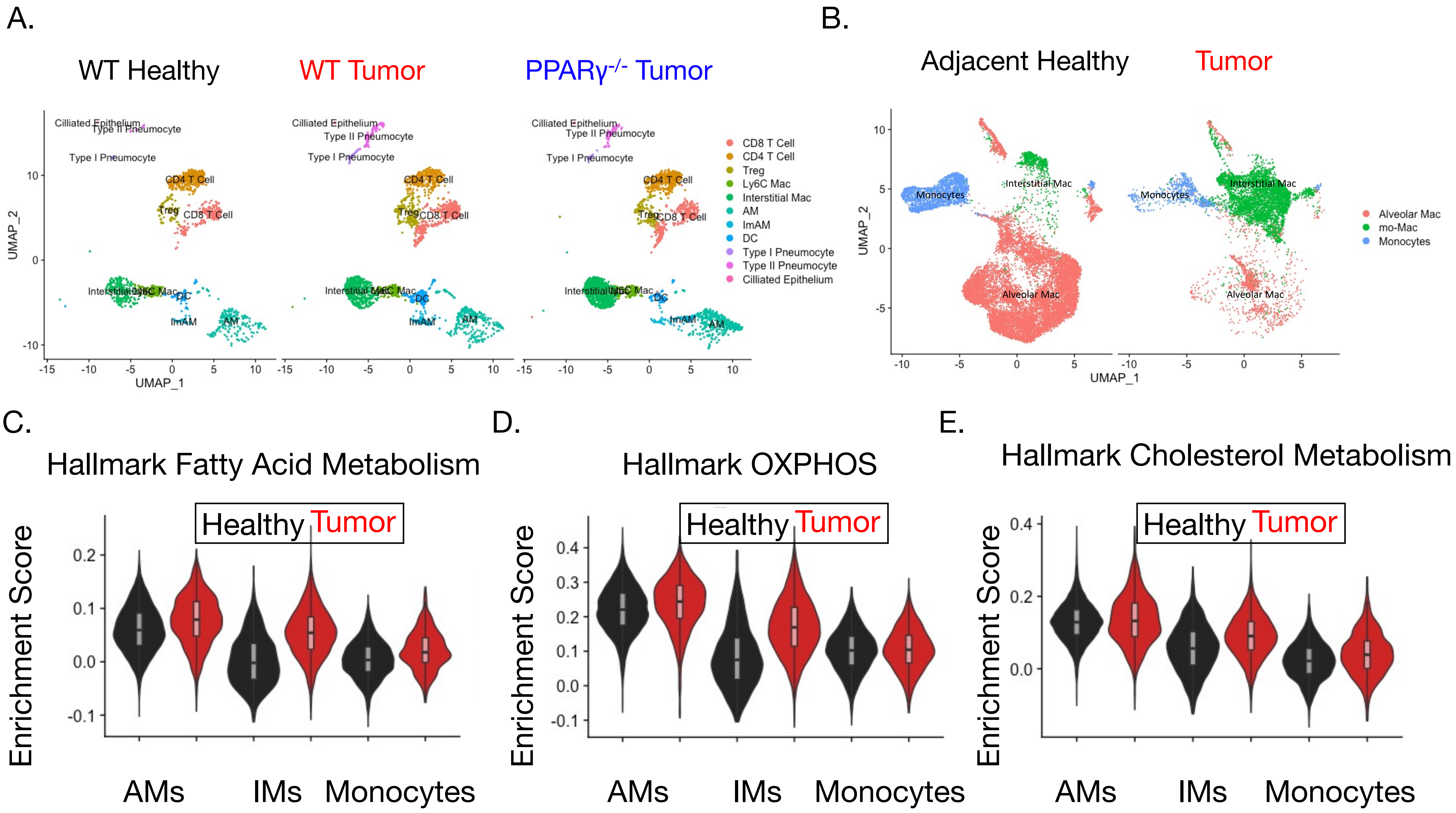


Kuhlmann et al., 2023
**Figure S4**

**Figure S4: Tumorigenesis rewires LUAD patient macrophage lipid and mitochondrial metabolism.**

(A) Annotated UMAP dimensional reduction of scRNA Seq from WT Healthy, WT Tumor and PPARγ$^{-/-}$ Tumors (CSF1R$^{Cre}$) based on canonical cell markers. (B) Annotated UMAP of myeloid cells in human LUAD and matched adjacent healthy tissue (data and cell types from GSE131907). (C-E) Violin plots show GSEA enrichment scores of hallmark pathway for (C) Fatty Acid Metabolism, (D) OXPHOS and (E) Cholesterol Metabolism in the different myeloid cells found in human LUAD or NSCLC tumors (red) and healthy adjacent tissue (black) using scRNA-seq datasets (data from GSE131907).

**Supplemental Table 1:**
**ST1:BALF Patient Sample Clinical Features**

| Sex | Age | Smoker | Cancer | Cancer Tissue | Other Complications |
|---|---|---|---|---|---|
| F | 52 | Y (current) | N | N/A | COPD |
| F | 76 | N | Y | Large B-cell lymphoma | COPD |
| M | 66 | Y (former) | Y | Squamous cell skin cancer | COPD |
| M | 59 | N | Y | papillary adenocarcinoma of thyroid | pulmonary disease due to mycobacteria |
| F | 75 | Y (former) | N | N | pulmonary disease due to mycobacteria |
| F | 58 | Y (former) | Y | LUAD | N/A |
| F | 68 | N | Y | Lung Cancer | Cholangiocarcinoma; type 2 diabetes; CKD (chronic kidney disease) |
| F | 87 | Y (former) | Y | Lung Cancer | pancreatic neoplasm |
| F | 64 | N | Y | LUAD | secondary malignant neoplasm to brain and spinal cord; bone mets |
| M | 67 | Y (former) | Y | LUAD | mets to brain, LN, liver, bone |
